# Dissecting human skeletal stem cell ontogeny by single-cell transcriptomic and functional analyses

**DOI:** 10.1101/2020.12.22.423948

**Authors:** Jian He, Jing Yan, Jianfang Wang, Liangyu Zhao, Qian Xin, Yang Zeng, Yuxi Sun, Han Zhang, Zhijie Bai, Zongcheng Li, Yanli Ni, Yandong Gong, Yunqiao Li, Han He, Zhilei Bian, Yu Lan, Chunyu Ma, Lihong Bian, Heng Zhu, Bing Liu, Rui Yue

## Abstract

Human skeletal stem cells (SSCs) have been discovered in fetal and adult bones. However, the spatiotemporal ontogeny of human SSCs during embryogenesis has been elusive. Here we map the transcriptional landscape of human embryonic skeletogenesis at single-cell resolution to address this fundamental question. We found remarkable heterogeneity within human limb bud mesenchyme and epithelium, as well as the earliest osteo-chondrogenic progenitors. Importantly, embryonic SSCs (eSSCs) were found in the perichondrium of human long bones, which self-renew and generate osteochondral lineage cells, but not adipocytes or hematopoietic stroma. eSSCs are marked by the adhesion molecule CADM1 and highly enrich FOXP1/2 transcriptional network. Interestingly, neural crest-derived cells with similar phenotypic markers and transcriptional network were also found in the sagittal suture of human embryonic calvaria. Taken together, this study revealed the cellular heterogeneity and lineage hierarchy during human embryonic skeletogenesis, and identified distinct skeletal stem/progenitor cells that orchestrate endochondral and intramembranous ossification.

## Introduction

Multipotent and self-renewing skeletal stem cells (SSCs) were discovered in the growth plate of early postnatal mice by phenotypic profiling and lineage tracing studies^1,2^. SSCs were also found within PTHrP^+^ chondrocytes in the resting zone of mouse postnatal growth plate^3^, as well as in the periosteum of postnatal long bones and calvaria (also known as periosteal stem cells, PSCs)^4^. Importantly, SSCs were recently identified in the growth plate of 17-week-old human long bones, suggesting that they are evolutionarily conserved in human fetus^5^. Similar to bone marrow stromal cells (BMSCs) that maintain the adult skeleton^6–9^, mouse and human SSCs from the growth plate give rise to chondrocytes, osteoblasts and hematopoietic stroma upon *in vivo* transplantation^1, 5^. However, they do not differentiate into adipocytes, highlighting the functional differences among SSCs at distinct developmental stages and anatomical sites^10, 11^. Whereas lineage tracing studies in mice revealed multiple waves of osteoprogenitors during skeletal development^12–14^, the embryonic origin of human SSCs during skeletogenesis remains unknown. Discovery of an evolutionarily conserved embryonic SSC population will not only clarify the spatiotemporal ontogeny of SSCs, but also shed light on novel cell therapies that promote skeletal regeneration.

In vertebrates, the earliest progenitors of appendicular skeleton are formed within limb buds^15, 16^. Limb patterning along the anterior-posterior (AP) axis is regulated by sonic hedgehog (SHH) signals from the zone of polarizing activity (ZPA)^17^, while the proximal-distal (PD) axis patterning is mainly regulated by FGF signals from the apical ectodermal ridge (AER)^18, 19^. The distal mesenchymal cells underlying AER are undifferentiated and highly proliferative when receiving the FGF and WNT signals^20, 21^, which form the progress zone that elongates the limb buds. The core mesenchyme outside progress zone express *SOX9* to specify the osteo-chondrogenic lineage and generate cartilage template. Although different mesenchymal progenitors have been identified in mouse and chick limb buds^22, 23^, the cellular heterogeneity and lineage hierarchy within human limb buds remain unknown.

After chondrogenic differentiation of limb bud mesenchymal progenitors, long bones are generated by endochondral ossification^24^. Blood vessels invade the center of cartilage template with perichondrial osteoprogenitors to form the primary ossification center (POC)^12, 14^, where osteoblasts, vascular endothelial cells, pericytes and hematopoietic cells populate to form the bone marrow^25–29^. In contrast to long bones, calvarial bones are generated by intramembranous ossification, which involves cranial mesenchyme condensation and direct mineralization on top of the cartilage anlagen^30–33^. Whereas long bones are derived from lateral plate mesoderm, calvarial bones are derived from both neural crest and paraxial mesoderm that generate different parts of the calvarium^34, 35^. Interestingly, although mouse long bone SSCs and calvarial PSCs are distinct stem cell populations that mediate endochondral and intramembranous ossification, respectively, they share similar phenotypic markers (Lineage^−^CD51^+/low^Thy1^−^6C3^−^CD200^+^CD105^−^)^1, 4^. Whether the embryonic long bones and calvaria contain skeletal stem/progenitor cells that share similar molecular features remain to be explored.

Single-cell RNA-sequencing (scRNA-seq) is a powerful tool in dissecting the cellular composition and lineage hierarchy within heterogeneous or rare cell populations^36–38^. In the musculoskeletal system, a high-throughput scRNA-seq study during mouse embryonic development reported the transcriptional landscapes of AER, limb bud mesenchyme and skeletal muscle before POC formation^39^. Recent scRNA-seq studies in adult mouse bone marrow also revealed the cellular heterogeneity of BMSCs, endothelial cells and osteo-chondrogenic lineage cells under homeostatic and stress conditions^40–42^. scRNA-seq profiling during axolotl limb regeneration identified convergence of connective tissue cells back to multipotent skeletal progenitors that formed a limb bud-like blastema structure^43^. In contrast, scRNA-seq studies in the human skeletal system are still lacking, especially during embryonic development.

In this study, we generated the first comprehensive human embryonic skeletogenesis cell atlas by scRNA-seq. By systematically examining the cellular heterogeneity and lineage hierarchies within multiple skeletal sites, we identified distinct skeletal stem/progenitor cells in human embryonic long bone and calvarium.

## Results

### Integrated analyses of single-cell transcriptomes during limb bud and long bone development

To test whether SSCs exist during embryogenesis, we analyzed human limb buds at 5 weeks post conception (5 WPC), as well as human limb long bones at 8 weeks post conception (8 WPC). Hematoxylin and eosin staining showed condensed mesenchyme within limb buds, and the nascent bone marrow cavity (POC) in the center of long bones (Fig. 1a). To map the single-cell transcriptomes, upper and lower limb buds (5 WPC, n=3, Supplementary information, Fig. S1a), as well as forelimb and hindlimb long bones (8 WPC, n=3, Supplementary information, Fig. S1a) were dissected and subjected to enzymatic digestions. Dissociated cells were then sorted by flow cytometry to obtain live single cells for 3’ scRNA-seq on a 10X Genomics platform (Fig. 1b). After quality control and doublet exclusion, we obtained 19,890 single cells in 5 WPC limb buds and 15,680 single cells in 8 WPC long bones (Supplementary information, Fig. S1a). On average, we detected 2,841 genes (10,212 unique molecular identities, UMI) per cell with less than 2.4% mitochondrial genes Supplementary information, Fig. S1a). Normal karyotype was inferred by calculating copy number variation (CNV) scores on 100 randomly sampled cells for each embryo (Supplementary information, Fig. S1b)^44^. We performed canonical correlation analysis (CCA) to normalize variance and correct batch effects among different samples^45^. Integrated analysis of the limb bud and long bone samples revealed 16 subsets (Fig. 1c and Supplementary information, Fig. S1c). The robustness of cell clustering was validated by calculating silhouette values (Supplementary information, Fig. S1d)^46^, and by random sampling and re-clustering analysis (Supplementary information, Fig. S1e).

We found three PRRX1^+^ mesenchymal subsets that mainly exist in 5 WPC limb buds (clusters 1-3), which differentially expressed *PDGFRA*, reflecting mesenchymal progenitors at different maturation stages (Fig. 1c-e)^22^. Notably, cluster 4 is a mesenchymal subset that equally distributed between limb bud and long bone samples, which expressed *PRRX1*, low level of *SOX9* and the highest level of *PDGFRA*, reminiscent of osteo-chondrogenic progenitors (OCPs) that give rise to long bones (Fig. 1c-e)^22^. EPCAM^+^ epithelial cells (clusters 14 and 15)^47^ and GYPA^+^ erythrocytes (cluster 13)^48^ were mainly detected in limb buds, while SIX1^+^ myoprogenitors (cluster 9)^49^, CDH5^+^ endothelial cells (cluster 11)^50^ and CD68^+^ macrophages (cluster 12)^51^ were found in both samples (Fig. 1c-e). In contrast, RUNX2^+^ osteoprogenitors (cluster 5)^52^, OSR2^+^NOV^+^ perichondrial mesenchymal stromal cells (PMSCs, cluster 6)^53, 54^, SOX9^+^ chondroblasts and chondrocytes (clusters 7 and 8)^55^, MYOG^+^ myocytes (cluster 10)^56^, as well as SOX10^+^ Schwann cells (cluster 16)^57^ were mainly detected in long bones (Fig. 1c-e and Supplementary information, Table S1).

**Fig. 1.**
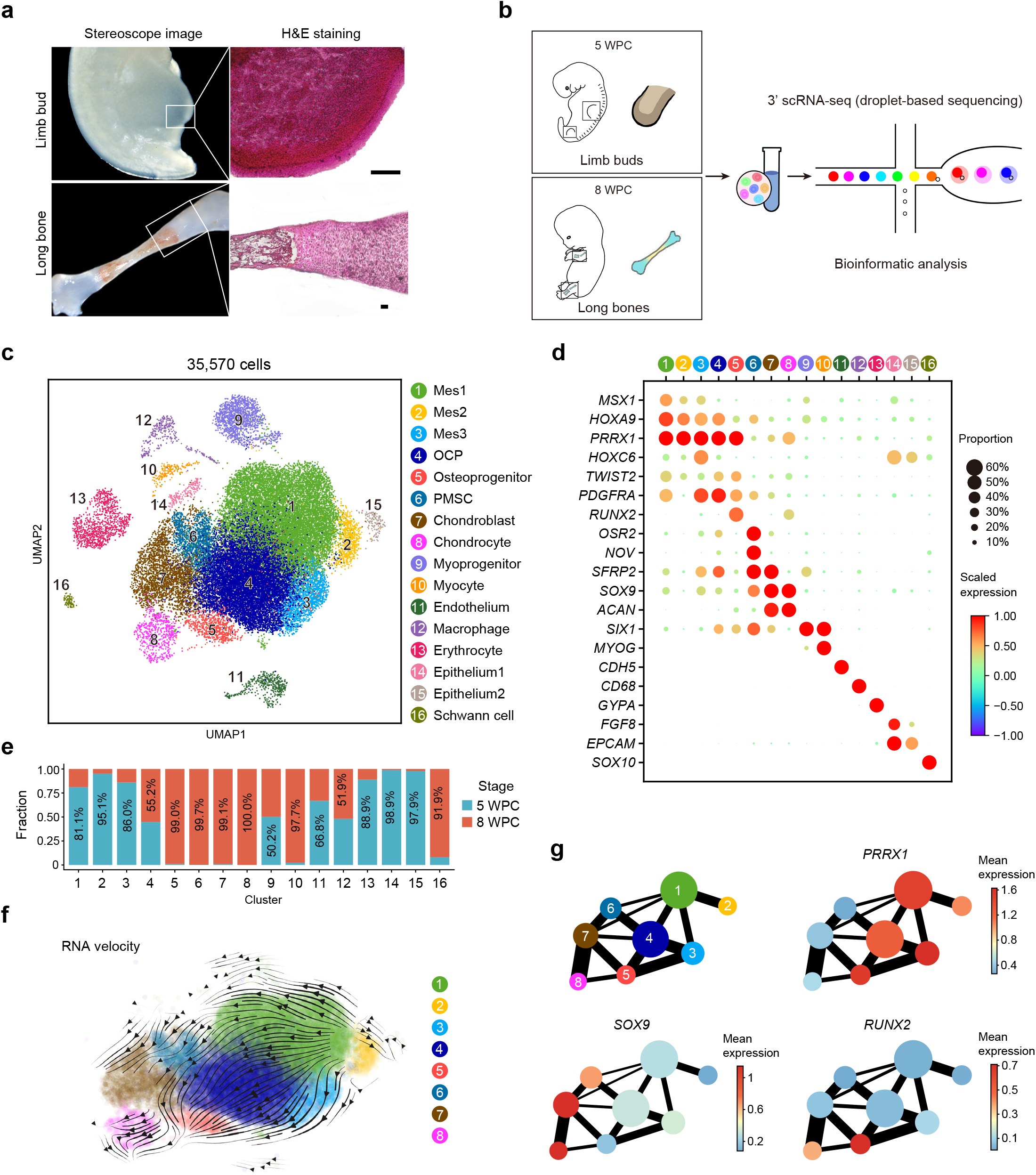
Integrated analysis of human limb buds and embryonic long bones. **a,** Representative stereoscope images (left) and H&E images (right) of 5 WPC human limb bud and 8 WPC human long bone. Scale bars: 100 μm. **b,** Sampling workflow and experimental scheme. Human embryonic cells from 5 WPC limb buds and 8 WPC long bones were sorted and subjected to droplet-based scRNA-seq. **c,** Distribution of 35,570 cells from limb buds and long bones. 16 subsets were visualized by uniform manifold approximation and projection (UMAP). **d,** Dot plots showing the expression of curated feature genes in 16 subsets. Dot size represented the proportion of cells expressing specific gene in the indicated subset and color bar represented the gene expression levels. **e,** Proportion of cells from 5 WPC limb buds and 8 WPC long bones in each subset. **f,** Developmental trajectory inferred by RNA velocity and visualized on the UMAP projection. **g,** Partition-based graph abstraction (PAGA) showing the connectivity among subsets in (**f**). The mean expression of representative genes (Mesenchymal: *PRRX1*; Chondrogenic: *SOX9*; Osteogenic: *RUNX2*) in each subset was showed in abstracted graph. Line thickness indicated the strength of connectivity. Color bar represents the gene expression levels.

Pearson correlation analysis clearly distinguished the skeletogenic and non-skeletogenic subsets (Supplementary information, Fig. S1f). Pseudotime analysis by RNA velocity^58^ showed a differentiation continuum from limb bud mesenchymal progenitors to OCPs, followed by cell fate specification into osteogenic and chondrogenic lineages (Fig. 1f). Partition-based graph abstraction (PAGA) analysis^59^ showed a pivotal role of OCPs in linking limb bud mesenchymal progenitors (PRRX1^+^) to PMSC/chondroblasts/chondrocytes (SOX9^+^) and osteoprogenitors (RUNX2^+^) in embryonic long bones (Fig. 1g). Next, we focused on this OCP lineage and separately analyzed the limb bud and long bone samples to trace back the origin of SSCs.

### Delineating mesenchymal lineage specification during limb bud development

We were able to identify 10 subsets in 5 WPC human limb buds (Fig. 2a). Hierarchical analysis within the 4 mesenchymal subsets showed that Mes1 (cluster 1) clustered with Mes2 (cluster 2), while Mes3 (cluster 3) and OCP (cluster 4) clustered together (Fig. 2b). Of the two epithelial subsets, only cluster 9 highly expressed AER marker *FGF8* (Fig. 1d), consistent with previous study in mouse embryos (Fig. 2b)^39^. Surprisingly, PAGA analysis found a strong correlation between Mes2 and epithelial subsets (Fig. 2c), raising the possibility that Mes2 might correspond to progress zone mesenchyme that lies underneath the limb bud epithelium^16, 60^. Consistent with this hypothesis, cell cycle analysis showed that Mes2 was more proliferative as compared to other mesenchymal subsets, with more cells in G2/M phase (Fig. 2d). Gene ontology (GO) analysis showed that Mes2 enriched genes regulating metabolic processes, while Mes3 and OCP enriched genes involved in embryonic skeletal development and ossification (Fig. 2e).

**Fig. 2.**
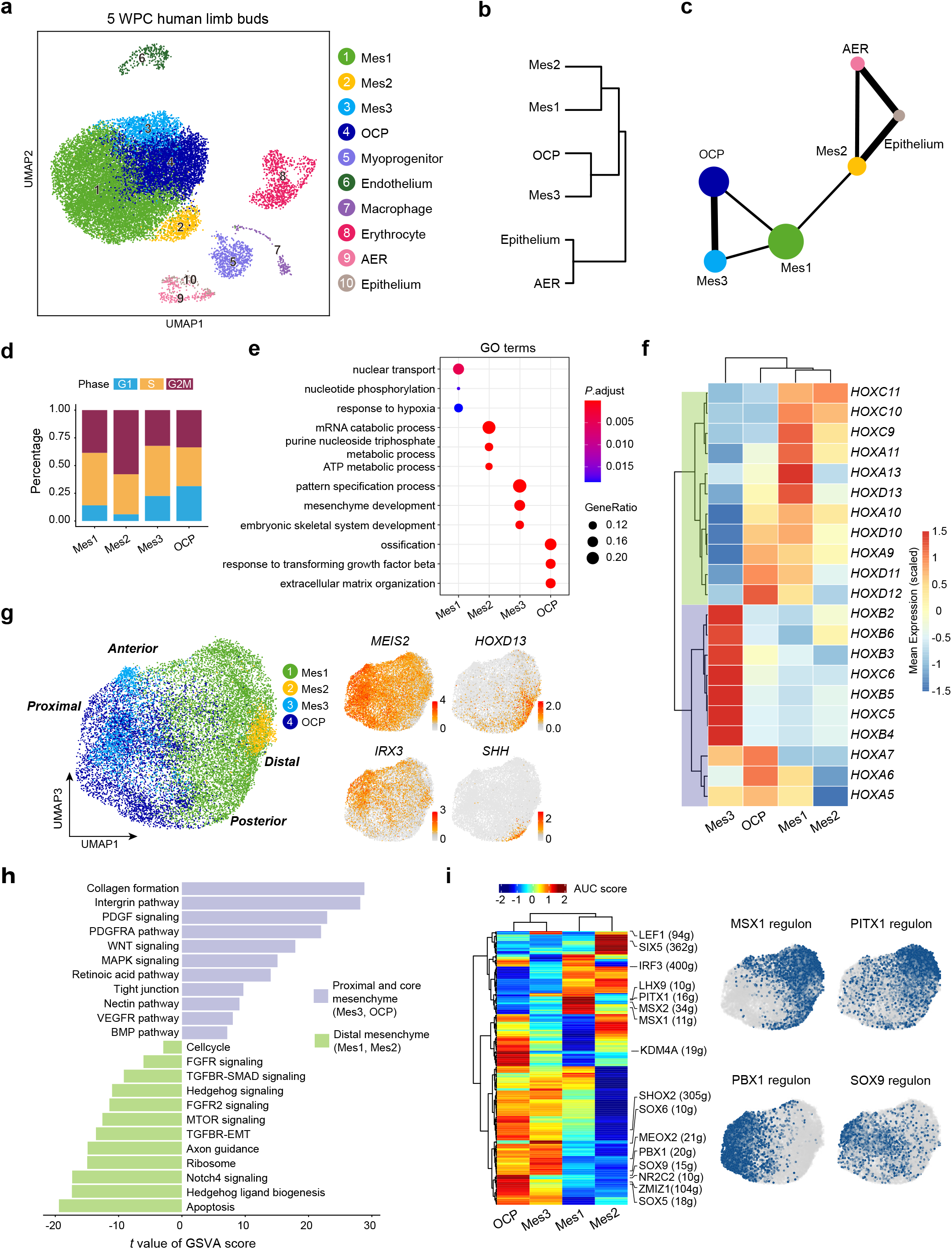
Characterization of human limb bud mesenchyme and epithelium. **a,** UAMP visualization of the 10 subsets in 5 WPC limb buds. **b,** Hierarchical clustering of the mesenchymal and epithelial subsets using top 50 principal components (PCs). **c,** The inferred relationships among the mesenchymal and epithelial subsets in PAGA layout. **d,** Stacked bar charts showing the cell cycle distributions in the mesenchymal subsets. **e,** Enriched GO terms of differentially expressed genes (DEGs) in the mesenchymal subsets. **f,** Heatmap showing expression of curated HOX genes scaled across the mesenchymal subsets. Hox genes were clustered into two branches based on hierarchical clustering of the rows, as indicated in green and purple. **g,** Visualization of the mesenchymal subsets (left) with UMAP plots showing the expression of curated PD and AP marker genes (right; Proximal: MEIS2; Distal: HOXD13; Anterior: IRX3; Posterior: SHH). **h,** GSVA analysis of pathway enrichment in the proximal and core mesenchyme (Mes3/OCP) and distal mesenchyme (Mes1/2). T values for each pathway were shown (two-sided unpaired limma-moderated t test). **i,** Heatmap showing the area under the curve (AUC) score of regulons enriched in the mesenchymal subsets. Z-score (row scaling) was computed. Hierarchical clustering on rows and columns indicated regulon patterns and correlation between cell subsets, respectively. AUC of representative regulons were shown by UMAP plots.

During limb bud outgrowth, *HOX* gene expressions switch from 3’ to 5’ topologically associating domains along the PD axis^61^. We found that Mes3 preferentially expressed 3’ *HOX* genes such as *HOX2-6*, while Mes1 and Mes2 preferentially expressed 5’ *HOX* genes such as *HOX9-11*, suggesting that they represented proximal (Mes3) and distal (Mes1 and Mes2) mesenchymal cells, respectively (Fig. 2f). In contrast, OCP expressed both 3’ and 5’ *HOX* genes, reminiscent of the core mesenchyme that gives rise to skeletal tissues (Fig. 2f). Consistent with this, when we aligned the mesenchymal subsets along PD and AP axes using known marker genes such as *MEIS2*, *IRX3*, *HOXD13* and *SHH* (Fig. 2g), Mes3 and OCP were positioned at the proximal end, while Mes1 and Mes2 were positioned at the distal end (Fig. 2g). Of note, the distal most localization of Mes2 was in line with the progress zone. Consistent with previous studies^62, 63^, gene set variation analysis (GSVA) showed that the proximal and core mesenchyme enriched genes related to retinoic acid and PDGF signaling, while the distal mesenchyme enriched genes related to Hedgehog, FGF, TGFβ and Notch signaling (Fig. 2h). To explore the gene regulatory networks (regulons) that determine cell fate specification in the mesenchymal subsets, we applied single-cell regulatory network inference and clustering (SCENIC) method to score the activity of regulons by an AUCell algorithm (AUC score), which reflects the co-expression of transcription factors (TFs) and their downstream target genes in each individual cell^64^. Hierarchical clustering of the AUC scores again distinguished proximal/core and distal mesenchymal subsets (Fig. 2i). MSX1 and PITX1 regulons were enriched in Mes1 and Mes2^65, 66^, while PBX1 and SOX9 regulons were enriched in Mes3 and OCP^22, 67^. Interestingly, we also identified several OCP-specific regulons such as ZMIZ1, NR2C2 and KDM4A, suggesting novel chondrogenic regulators within the limb bud mesenchyme (Fig. 2i and Supplementary information, Table S2).

To explore evolutionarily conserved and species-specific features during limb bud development, we analyzed a recently published scRNA-seq dataset of mouse hindlimb buds at similar embryonic stage (E11.5) (Supplementary information, Fig. S2a)^68^. SciBet is a recently developed algorithm that predicts cell identity by training multinomial model with given dataset^69^. By training SciBet with our human dataset, we found that most human subsets were conserved in mouse except that Mes2 and epithelium (non-AER) subsets were not predicted in mouse limb buds (Supplementary information, Fig. S2b and Table S1). The lack of a highly proliferative Mes2 subset implied advanced maturation of E11.5 mouse limb buds (Supplementary information, Fig. S2a)^70^. Consistent with this, mouse OCP subset highly expressed *SOX9* (Supplementary information, Fig. S2c), suggesting early chondrogenic differentiation. A much lower proportion of mouse AER was found within limb bud epithelium (6%) as compared to human AER (69%, Supplementary information, Fig. S2d), which could possibly explain why mouse limbs are much shorter than human limbs.

Taken together, these data revealed the cellular heterogeneity and species-specific features of human limb bud mesenchyme and epithelium. Since osteogenesis is not initiated in 5 WPC human limb buds, we went on to analyze the 8 WPC human long bones in search of embryonic SSCs.

### Delineating osteochondral lineage specification during long bone development

We analyzed the long bone dataset from 8 WPC human embryos (Supplementary information, Fig. S3a) and divided the osteochondral lineage cells (OCLCs) into 7 subsets (Fig. 3a). In addition to previously identified osteoprogenitor, PMSC, chondroblast and chondrocyte subsets (Fig. 3a, clusters 4-7), long bone OCPs (Fig. 1c) were subdivided into 3 subsets (clusters 1-3). Cluster 1 highly expressed *CXCL12* and *PDGFRA* (Supplementary information, Fig. S3b and Table S1), which are markers of BMSCs^28, 71^. Cluster 2 highly expressed *TWIST2* that functions as an inhibitor of osteoblastic differentiation^72^, reminiscent of OCPs that were derived from limb bud mesenchyme. Cluster 3 highly expressed *GAS2*, *PTN* and localized in the center of all OCLC subsets (Fig. 3a, Supplementary information, Fig. S3a,b and Table S1). GO analysis showed significant enrichment of genes related to organ and appendage morphogenesis in clusters 1-3 (Fig. 3b). Interestingly, genes related to stem cell proliferation were enriched in cluster 3 (Fig. 3b), suggesting it might contain embryonic SSCs (eSSCs).

**Fig. 3.**
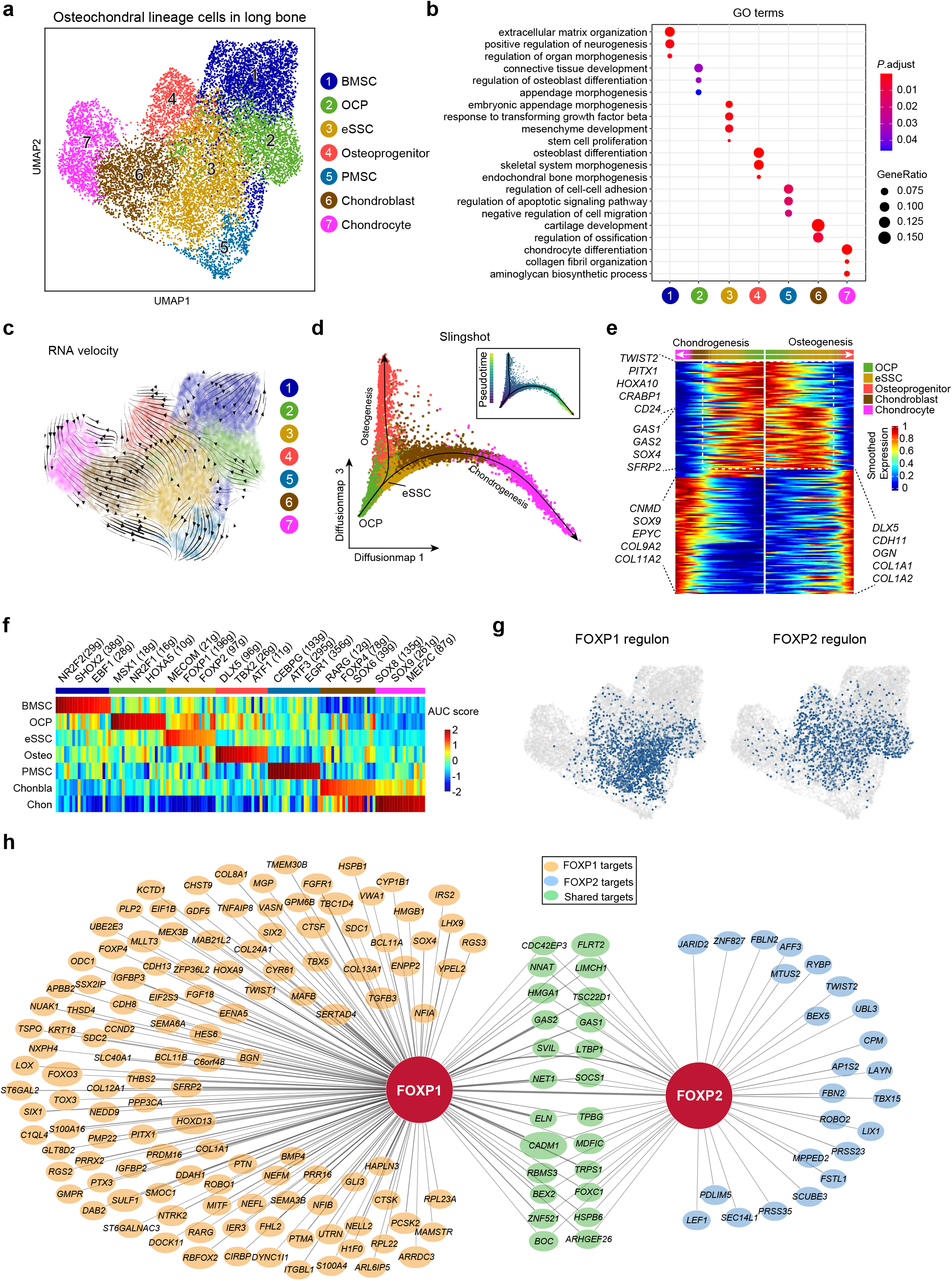
Characterization of the osteochondral lineage in human long bones identified embryonic SSCs. **a,** UMAP visualization of 7 OCLC subsets in 8 WPC human long bones. **b,** Enriched GO terms of differentially expressed genes (DEGs) among the 7 OCLC subsets. **c,** Developmental trajectory of 7 OCLC subsets inferred by RNA velocity and visualized on the UMAP projection. **d,** UMAP visualization of the osteogenic and chondrogenic trajectories simulated by Slingshot across OCP, eSSC, osteoprogenitor, chondroblast and chondrocyte subsets. The corresponding diffusion pseudotime was indicated in the upper right frame. **e,** Heatmap of gene expressions (smoothed over 20 adjacent cells) in OCP, eSSC, osteoprogenitor, chondroblast and chondrocyte subsets ordered by pseudotime of osteogenesis and chondrogenesis in (**d**). Top 200 genes were selected according to the *P* values of GVM test and representative genes were shown. Shared genes in the two trajectories were indicated in dashed box. **f,** Heatmap showing the AUC score of regulons enriched in human OCLC subsets. Z-score (column scaling) was calculated. Representative regulons were shown on the top. The number of predicted target genes for each regulon was shown in the parenthesis. **g,** AUC of FOXP1 and FOXP2 regulons were shown by UMAP plots. **h,** The FOXP1 and FOXP2 regulon networks in OCLC subsets. Line thickness indicated the level of GENIE3 weights. Dot size indicated the number of enriched TF motifs.

To test this hypothesis *in silico*, pseudotime analysis by RNA velocity was performed to explore the lineage relationships among OCLC subsets (Fig. 3c). We observed strong directional streams from eSSC toward osteoprogenitor, chondroblast/chondrocyte and PMSC subsets (Fig. 3c). Interestingly, OCP was upstream of both eSSC and BMSC, which formed two differentiation trajectories to generate the skeleton and bone marrow stroma, respectively (Fig. 3c). Diffusion map analysis of OCP, eSSC, chondroblast/chondrocyte and osteoprogenitor subsets simulated two differentiation trajectories featuring chondrogenesis and osteogenesis (Fig. 3d). Consistent with the RNA velocity analysis, eSSC was located at the branching point of osteogenesis and chondrogenesis (Fig. 3d). We set OCP as the root to identify temporally expressed genes over pseudotime, and found that genes highly expressed in OCPs (eg. *PITX1*, *HOXA10*, *CRABP1*, *CD24*) and eSSCs (eg. *GAS1/2*, *SOX4* and *SFRP2*) were gradually down-regulated, while genes that highly expressed in chondrocytes (eg. *CNMD*, *EPYC*, *COL9A2*, *COL11A2*) and osteoprogenitors (eg. *DLX5*, *CDH11*, *OGN* and *COL1A1/2*) were up-regulated upon terminal differentiation (Fig. 3e). SCENIC analysis showed that eSSCs highly enriched regulons such as FOXP1 and FOXP2 (Fig. 3f and Supplementary information, Table S2). The FOXP1 regulon seemed to be more specific to eSSCs, as the FOXP2 regulon was also enriched in OCPs and osteoprogenitors (Fig. 3g). Nevertheless, FOXP1/2 did share a significant amount of target genes in eSSCs (Fig. 3h).

We also analyzed a published scRNA-seq dataset of mouse hindlimb long bones at similar embryonic stage (E15.5) (Supplementary information, Fig. S3c,d)^68^. SciBet analysis found that human eSSC was evolutionarily conserved in mouse long bones (Supplementary information, Fig. S3e). Interestingly, FOXP1/2/4 regulons were highly enriched in mouse eSSCs (Supplementary information, Fig. S3f,g and Table S2), suggesting a fundamental role of FOXP family TFs in regulating eSSC specification. Taken together, we identified an eSSC subset among OCPs that could potentially regulate long bone development and POC formation.

### Identification of CADM1 as a phenotypic marker of eSSC

To prospectively isolate eSSCs for functional validation *ex vivo*, we first screened for cell surface markers that were differentially expressed among long bone OCLC subsets. Interestingly, we found the cell adhesion molecule *CADM1* to be preferentially expressed in eSSCs (Fig. 4a). SCENIC analysis showed that FOXP1/2 binding motifs were highly enriched in the predicted cis-regulatory elements of *CADM1* among all co-expressed target genes (Fig. 3h), suggesting that it could be used as a legitimate phenotypic marker of eSSCs. Since *CADM1* was also expressed in Schwann cells (Fig. 4a), we sought to further enrich eSSCs by combining with previously reported SSC and BMSC markers (Fig. 4a)^1, 5, 11^, and found that *PDPN* was differentially expressed in eSSCs (PDPN^+^) and Schwann cells (PDPN^−^) (Fig. 4a). Immunostaining of CADM1 and PDPN on 8 WPC human long bone sections showed that PDPN^+^CADM1^+^ cells mainly localize in the perichondrium surrounding POC and articular surface (Fig. 4b and Supplementary information, Fig. S4b), indicating their ability to generate chondrocytes and PMSCs. A few PDPN^+^CADM1^+^ cells were also found inside POC (Fig. 4b), reminiscent of osteoprogenitors that invade the cartilage template^12^.

**Fig. 4.**
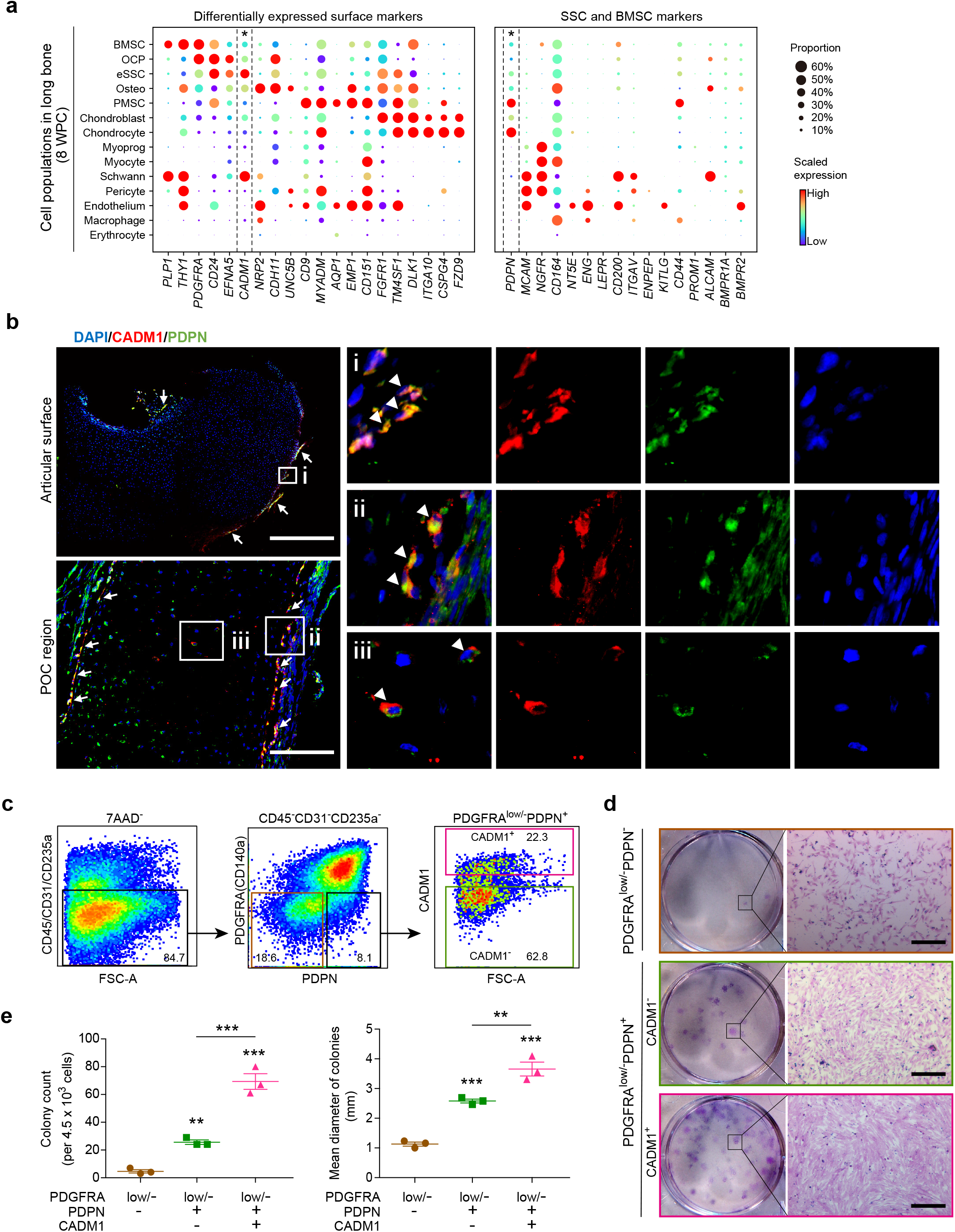
Identification of CADM1 as a phenotypic marker of eSSCs. **a,** Dot plots showing the expression of differentially expressed cell surface genes (left) and candidate SSC markers (right) in 8 WPC human long bone subsets. Asterisks indicated positive markers that were used to enrich eSSCs. **b,** Immunofluorescent images of PDPN^+^CADM1^+^ cells in 8 WPC human long bones. Overviews of PDPN^+^CADM1^+^ cells (arrows) in the articular (upper left) and POC (bottom left) regions were shown on the left. PDPN^+^CADM1^+^ cells were found in the inner layer of perichondrium in articular regions (i) and surrounding POC (ii). A few PDPN^+^CADM1^+^ cells were also found inside POC (iii). Arrow heads indicated enlarged PDPN^+^CADM1^+^ cells. Merged and single-channel images of DAPI (blue), CADM1 (red) and PDPN (green) were shown. Scale bars: 200 μm. **c,** Flow cytometry gating strategies for sorting different populations in 8 WPC long bones. **d,** Representative crystal violet staining of CFU-F colonies generated by the sorted populations as indicated in (**c**). Magnified images of the boxed areas were shown on the right. Scale bars: 25 μm. **e,** Quantifications of the number (top) and mean diameter (bottom) of the CFU-F colonies. The statistical significance of differences was determined using one-way ANOVA with multiple comparison tests (LSD). * *P* < 0.05; ** *P* < 0.01; *** *P* < 0.001. Error bars indicated SEM.

*In silico* transcript-averaged cell scoring (TACS) analysis^73^ revealed that the purity of eSSCs could be further enriched by PDGFRA^low/-^PDPN^+^CADM1^+^ cells among OCLC subsets (Supplementary information, Fig. S4a). In contrast, *THY1* (CD90), *NGFR* (CD271), *MCAM* (CD146) or *NT5E* (CD73) were hardly detected in eSSCs (Fig. 4a and Supplementary information, Fig. S4a). Next, we sorted PDGFRA^low/-^PDPN^−^, PDGFRA^low/-^PDPN^+^CADM1^−^and PDGFRA^low/-^PDPN^+^CADM1^+^ cells from 8 WPC human long bones by flow cytometry (Fig. 4c), and performed colony-forming unit-fibroblast (CFU-F) and mesenchymal sphere cultures to assess their colony-and sphere-forming efficiencies *ex vivo*. As compared to PDGFRA^low/-^PDPN^−^cells, PDGFRA^low/-^PDPN^+^CADM1^−^cells showed significantly increased colony-forming efficiency with colonies of larger size (Fig. 4d,e). Remarkably, PDGFRA^low/-^PDPN^+^CADM1^+^ cells showed an even higher colony-forming efficiency with significantly more colonies of larger size as compared to PDGFRA^low/-^PDPN^−^and PDGFRA^low/-^PDPN^+^CADM1^−^cells (Fig. 4d,e). Mesenchymal sphere formation analysis showed similar results (Supplementary information, Fig. S4c,d), suggesting that eSSCs highly enrich clonogenic activity.

### eSSCs self-renew and undergo osteo-chondrogenic differentiation

To test the self-renewal and differentiation potentials of eSSCs, we sorted PDGFRA^low/-^PDPN^+^CADM1^+^ cells to perform serial CFU-F colony formation assay, as well as trilineage differentiation (adipogenic, osteogenic and chondrogenic) both *in vitro* and *in vivo*. Single CFU-F colonies formed by flow cytometrically sorted PDGFRA^low/-^PDPN^+^CADM1^+^ cells were clonally expanded and serially passaged, which could generate secondary and tertiary colonies that maintain eSSC immunophenotypes (Fig. 5a and Supplementary information, Fig. S5a). Next, we performed *in vitro* trilineage differentiation of nonclonal and clonal cultures (cells were clonally expanded from single CFU-F colonies) of PDGFRA^low/-^PDPN^+^CADM1^+^ cells, and found that they underwent osteogenic and chondrogenic differentiation, but not adipogenic differentiation (Fig. 5b, and Supplementary information, Fig. S5b,c). The differentiation efficiency was quantified by qPCR analysis of adipogenic (*ADIPOQ* and *PPARG*), osteogenic (*RUNX2* and *SP7*) and chondrogenic (*SOX9* and *COL2A1*) marker genes (Fig. 5c and Supplementary information, Fig. S5d).

**Fig. 5.**
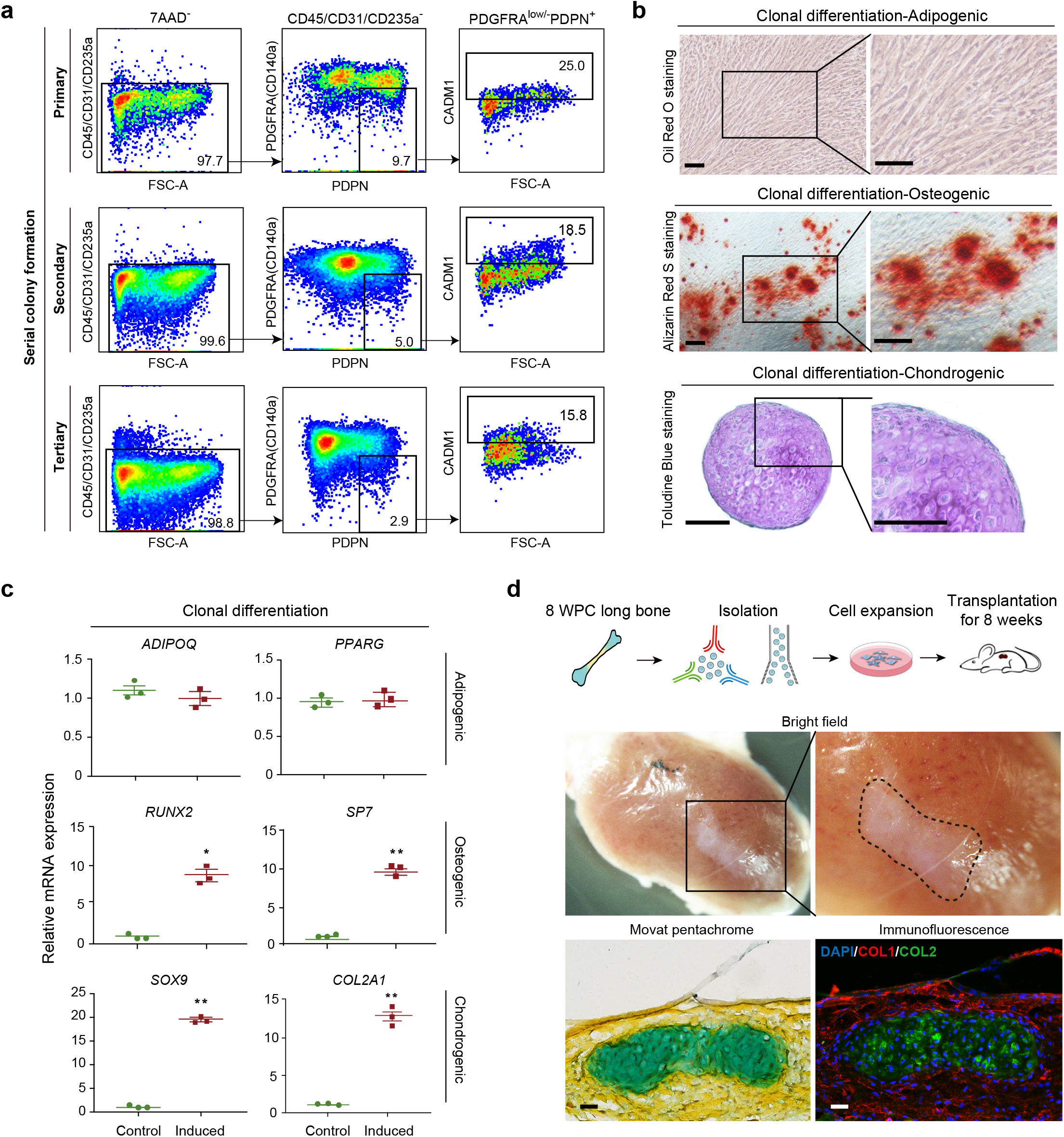
Functional characterizations of eSSCs *in vitro* and *in vivo*. **a,** Flow cytometry plots showing the maintenance of phenotypic eSSCs after serially passaging clonally expanded PDGFRA^low/-^PDPN^+^CADM1^+^ cells. **b,** Representative oil red O (top), alizarin red (middle) and toluidine blue (bottom) staining after adipogenic, osteogenic and chondrogenic differentiation of clonally expanded eSSCs (PDGFRA^low/-^PDPN^+^CADM1^+^). Magnified images of the boxed areas were shown on the right. Scale bars: 200 μm. **c,** qPCR analyses of adipogenic, osteogenic and chondrogenic marker genes in clonally expanded eSSCs before and after trilineage differentiation *in vitro*. The statistical significance of differences was determined using Wilcoxon signed rank test. * *P* < 0.05; ** *P* < 0.01. Error bars indicated SEM. **d,** Renal subcapsular transplantation. The work flow for functional characterization of eSSC *in vivo* (top). Subcapsular xenografts were dissected and sectioned 8 weeks after transplantation of culture expanded eSSCs into immunodeficient mice. Bright field (middle), Movat pentachrome staining (bottom left, cartilage: blue, bone and fibrous tissue: yellow) and immunofluorescent staining images (bottom right, DAPI: blue, collagen I: red, collagen II: green) were shown. Scale bars: 50 μm.

To test the differentiation potential of eSSCs *in vivo*, we performed renal subcapsular transplantation of cultured PDGFRA^low/-^PDPN^+^CADM1^+^ cells in immunodeficient mice. Eight weeks after transplantation, the subcapsular grafts were harvested and sectioned. Movat pentachrome staining and immunofluorescent staining of collagen I and II revealed osteo-chondrogenic differentiation of eSSCs (Fig. 5d). We did not observe bone marrow formation in the subcapsular grafts, suggesting that eSSCs are functionally distinct from growth plate SSCs that could organize a hematopoietic microenvironment^5^. Taken together, these data suggested that CADM1 is an important phenotypic marker of eSSCs, and that PDGFRA^low/-^PDPN^+^CADM1^+^ cells enriched self-renewing eSSCs that generate the osteochondral lineages during long bone development.

### Delineating osteogenic lineage specification during calvaria development

To test whether similar skeletal stem/progenitor cells exist in embryonic calvarium, we performed scRNA-seq in 8 WPC human calvaria (n=2, Supplementary information, Fig. S6a). Analysis of 7,287 CD235A^−^7AAD^−^(live non-erythrocytes) single cells revealed 12 distinct subsets (Fig. 6a), which included: 1) NGFR^+^ cranial neural crest (NC) cells (cluster 1) that highly expressed *NES*^74^; 2) Two GJA1^+^ subsets including vascular leptomeningeal cells (cluster 2, VLMCs) that highly expressed *SLC6A13* and *PTGDS*^75^, and migratory NC (mig_NC) cells that expressed higher level of *BMP4* (cluster 3)^76, 77^; 3) Neural crest-derived cells (cluster 4, NCDC) that highly expressed *BMP4* and *FOXC2*^78^; 4) RUNX2^+^ osteoprogenitors (cluster 5) that highly expressed osteogenic factors *DLX5* and *CLEC11A*^79, 80^; 5) Two OSR2^+^ PMSC subsets (clusters 6 and 7) that highly expressed *POSTN*; 6) SOX9^+^ chondrocytes that highly expressed *COL9A2* (cluster 8); 7) PDGFRB^+^ pericytes that highly expressed *MCAM* and *ACTA2*; 8) MYF5^+^ myoblasts; 9) CDH5^+^ endothelial cells and 10) CD68^+^ macrophages (Fig. 6a,b, Supplementary information, Table S1).

**Fig. 6.**
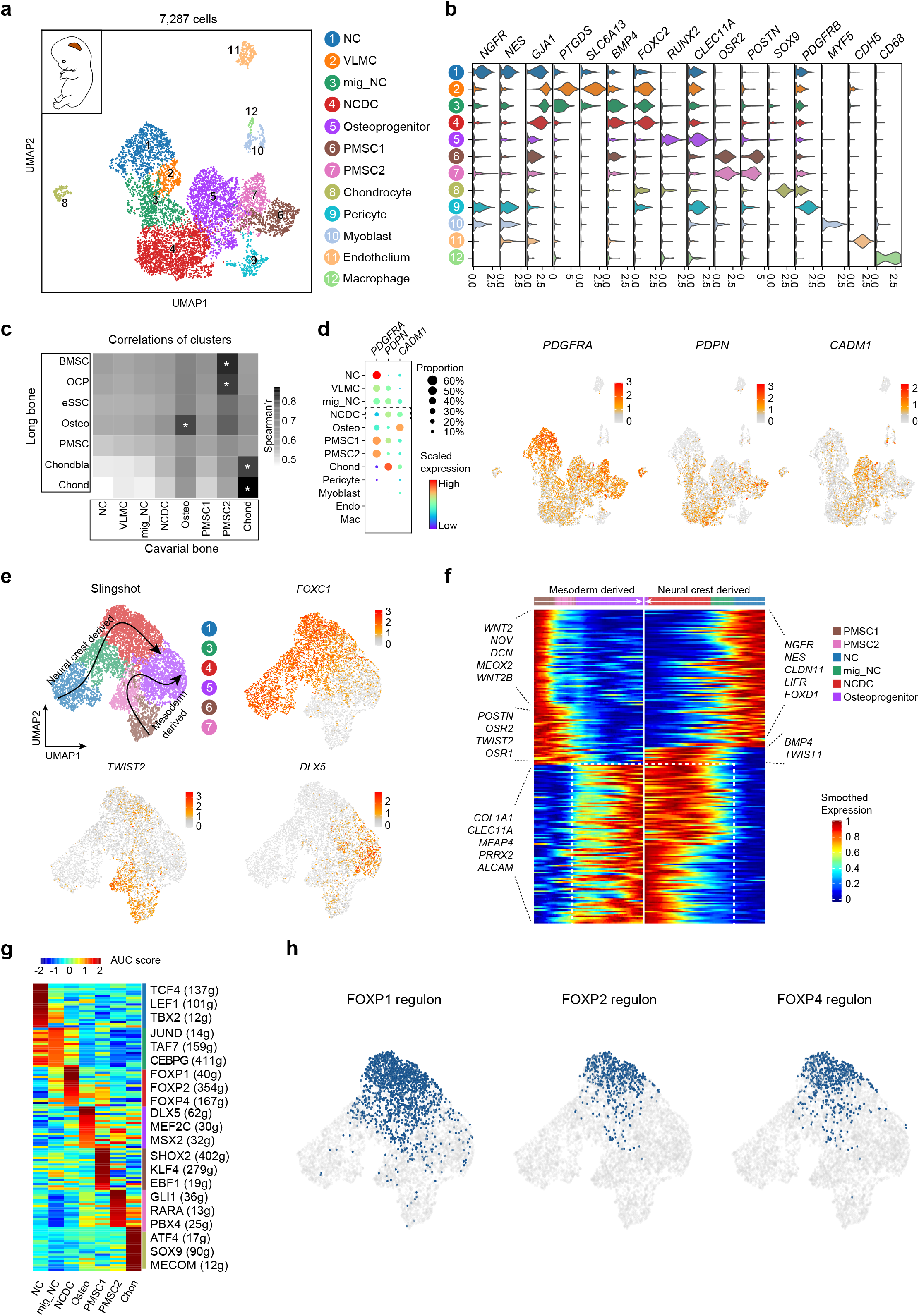
Characterization of the osteogenic lineages in human embryonic calvaria identified neural crest-derived skeletal progenitors. **a,** UMAP visualization of 12 subsets in 8 WPC calvarial bones. Inset illustrated the position of calvarial bone. **b,** Violin plots showing the expression of feature genes for each subset. **c,** Heatmap showing the transcriptome correlation between osteogenic subsets in calvarial and OCLC subsets in long bone. Asterisks indicated subsets with correlation coefficients > 0.8. **d,** Dot plots (left) and UMAP plots (right) showing the expression of eSSC marker genes subsets of 8 WPC calvarial. **e,** UMAP visualization of the two osteogenic trajectories simulated by Slingshot across NC, mig_NC, NCDC, osteoprogenitor, PMSC1 and PMSC2 subsets (Upper left). Expression UMAP plots of marker genes (NC: *FOXC1*; Mesoderm: *TWIST2*; Osteoprogenitor: *DLX5*). **f,** Heatmap of the gene expressions (smoothed over 20 adjacent cells) in subsets ordered by pseudotime of osteogenesis as in (**e**). Top 200 genes were selected according to the *P* values of GVM test and representative genes were shown. Shared genes in two trajectories were indicated in dashed box. **g,** Heatmap showing the AUC scores of regulons enriched in the osteogenic subsets. Z-score (row scaling) was computed. Representative regulons were shown on the right. **h,** AUC of FOXP1/2/4 regulons were shown by UMAP plots.

As compared to 8 WPC long bones, higher proportion of osteoprogenitors and PMSCs but much lower proportion of chondrocytes were detected in 8 WPC calvarial bones (Supplementary information, Fig. S6b), highlighting the fundamental differences between endochondral and intramembranous ossification^15^. Spearman correlation analysis showed that calvarial chondrocyte and osteoprogenitor subsets were more corelated with their long bone counterparts (Fig. 6c), while the PMSC2 subset seemed to be closely related to OCP and BMSC subsets in long bones (Fig. 6c). Integrated analysis of all subsets at the pseudo-bulk level showed similar results (Supplementary information, Fig. S6c). Although no calvarial subset highly resembled long bone eSSC at the transcriptome level, we did notice that NCDC shared similar phenotypic markers as long bone eSSC (PDGFRA^low/-^PDPN^+^CADM1^+^) (Fig. 6d). Immunostaining on 8 WPC human calvarial sections showed PDPN^+^CADM1^+^ cells in the outer layer of sagittal suture (Supplementary information, Fig. S6d), reminiscent of PSCs in adult mouse calvarium^4^.

To predict the functional role of NCDC during calvarial bone development, we performed pseudotime analysis within osteogenic subsets by Slingshot^81^, which revealed two distinct differentiation trajectories (Fig. 6e). Specifically, the FOXC1^+^ NC lineage cells and TWIST2^+^ mesodermal lineage cells converge to generate DLX5^+^ osteoprogenitors (Fig. 6e), where NCDC seemed to play a pivotal role in the transition from migratory NC cells to osteoprogenitors (Fig. 6e). Gene expression analysis showed that NC lineage cells down-regulated neural genes such as *NGFR, NES* and *CLDN11*^82^ to generate NCDCs and osteoprogenitors (Fig. 6f). In contrast, mesodermal lineage cells down-regulated WNT signaling genes such as *WNT2* and *WNT2B*, as well as TFs like *MEOX2*, *OSR1* and *OSR2* to generate osteoprogenitors (Fig. 6f). Calvarial osteoprogenitors highly expressed *COL1A1*, *PRRX2* and *CLEC11A*, a recently identified osteogenic factor that promotes the maintenance of adult skeleton^80, 83^. GSVA analysis showed that EPH-EPHRIN, WNT-LPR6 and RAC1 activation pathways were enriched in NCDCs (Supplementary information, Fig. S6e). Similar to long bone eSSC, SCENIC analysis showed that FOXP1/2 regulons were highly enriched in NCDC (Fig. 6g,h, Supplementary information, Fig. S6g and Table S2), although little FOXP1/2 target genes were shared by these two subsets (Fig. 3h and Supplementary information, Fig. S6f). In addition, the FOXP4 regulon was also enriched in NCDC and formed an integrated transcriptional network with FOXP1/2, suggesting a fundamental role of FOXP family TFs in NCDC specification. Taken together, these data revealed two distinct routes of osteogenic differentiation in calvaria, and identified NCDC as a potential skeletal stem/progenitor cell subset that mediates intramembranous ossification during calvarial development.

## Discussion

Whereas skeletogenesis has been extensively studied in model organisms such as mouse, chick and axolotl^39, 43, 68, 84^, human studies largely remain at the histomorphological level. In 2018, Ferguson et al. interrogated 17 WPC human fetal musculoskeletal subsets by bulk RNA-seq and compared chondrocyte features among 4 developmental stages^85^. Recently, a human skeletal muscle atlas was reported during embryonic, fetal and postnatal development^86^. Here, we provide the first transcriptional landscape of human embryonic skeletogenesis at single-cell resolution and shed light on novel skeletal stem/progenitor cells orchestrating lineage specifications during endochondral and intramembranous ossification. Together with the previous studies, we are now approaching a better understanding of the ontogeny of human musculoskeletal system.

Human SSCs were originally found in fetal long bones, which could be prospectively isolated by a combination of phenotypic markers (Lin^−^PDPN^+^CD146^−^CD73^+^CD164^+^)^5^. To test whether human SSCs exist during embryonic development, we mapped the single-cell transcriptomes in 5 WPC human limb buds and 8 WPC embryonic long bones and found an OCP subset that tightly links limb bud mesenchyme to endochondral ossification (Fig. 1f,g). Unlike mouse limb bud mesenchymal progenitors (Sox9^−^Pdgfra^hi^) and OCPs (Sox9^+^Pdgfra^hi^)^22^, human OCPs are SOX9^low^PDGFRA^hi^ (Fig. 1d), suggesting that they are less differentiated than mouse OCPs. We then focused on OCPs in both limb buds and long bones in order to identify skeletal stem/progenitor cells during human embryonic limb development.

Although the patterning mechanisms during limb bud development have been well-studied and simulated by different models^15, 17, 19^, the heterogeneity of human limb buds has been elusive. We identified 4 mesenchymal and 2 epithelial subsets in 5 WPC human limb buds. By analyzing *Hox* gene expression and well-known marker genes, we were able to align the 4 mesenchymal subsets along PD and AP axes (Fig. 2f,g). Importantly, we identified a highly proliferative Mes2 subset at the distal most mesenchyme, implicating immature mesenchymal progenitors underlying AER^21, 65^. We also identified an OCP subset with chondrogenic potential in the core mesenchyme. As compared to human limb buds, the E11.5 mouse limb buds lacked an equivalent Mes2 subset, showed early chondrogenic differentiation of OCP, and contained fewer proportion of AER cells (Supplementary information, Fig. S2). Together, these data suggested greater potential of human limb bud outgrowth that could possibly contribute to longer limb bones. Whether the novel regulons identified in human limb bud OCP (eg. ZMIZ1, NR2C2 and KDM4A) critically control chondrogenic differentiation remains to be validated by functional studies.

The OCPs in 8 WPC human long bones could be subdivided into 3 subsets, namely, OCP, BMSC and eSSC. The long bone OCP subset could be derived from limb bud OCPs, which generate BMSCs and eSSCs to form the bone marrow stroma and appendicular skeleton, respectively (Fig. 3c). Similar to human SSCs^5^, eSSCs were predicted to generate chondroblasts/chondrocytes, osteoprogenitors and PMSCs in 8 WPC long bones (Fig. 3c). Interestingly, PAGA analysis of integrated samples revealed a critical role of PMSC in mediating chondrogenic differentiation (Fig. 1g), which was not reflected by RNA velocity analysis in long bones (Fig. 3c). This discrepancy could be explained by the fact that RNA velocity analysis is more suitable for predicting differentiation trajectories in full-length sequencing dataset^87^. Since PMSC expressed higher level of SOX9 as compared to eSSC (Supplementary information, Fig. S3b), we tend to believe that PMSC also contributes to chondrogenic differentiation. Importantly, an equivalent eSSC subset was also found in E15.5 mouse embryonic long bones (Supplementary information, Fig. S3), suggesting its evolutionary conservation. The fact that both human and mouse eSSCs enriched FOXP1/2 regulons was quite intriguing (Fig. 3i), since mouse Foxp1/2/4 have been previously shown to regulate endochondral ossification by promoting chondrocyte proliferation and inhibiting osteoblast differentiation^88^. They do so by interacting with Runx2 to repress its transcriptional activity^88^, which could possibly explain how eSSCs are maintained in an undifferentiated state. Notably, much more FOXP2 target genes were found in human long bones as compared to mouse (human: 97, mouse: 12), consistent with a recent discovery that skeletal FOXP2 contributes to the acquisition of important human traits such as language and bipedalism^89^. More functional studies are needed to fully address the molecular mechanisms by which FOXP1/2 regulate human eSSC self-renewal and differentiation.

CADM1 was previously identified as an osteoblastic adhesion molecule and a diagnostic marker for osteosarcoma^90^. Here we found that PDPN^+^CADM1^+^ cells enriched eSSCs in 8 WPC human long bones, which mainly localize in the perichondrium surrounding POC and articular surface (Supplementary information, Fig. S4b). Interestingly, the perichondrial localization of eSSC was consistent with the expression pattern of Foxp1/2/4 in E13.5 mouse perichondrium^88^. A few PDPN^+^CADM1^+^ cells were also found inside the developing POC, which might represent invading osteoprogenitors derived from eSSCs^12^. Similar to human SSCs, eSSCs exhibit high clonogenic capacity, which self-renew and undergo osteochondral but not adipogenic differentiation *in vitro* and *in vivo*^5^. Notably, eSSCs do not form bone marrow upon renal subcapsular transplantation, suggesting that skeletogenic and hematopoietic functions might be segregated between eSSCs and BMSCs (Fig. 3c). However, whether the BMSC subset could support hematopoiesis in 8 WPC human embryo is still elusive, since fetal liver is the primary hematopoietic site at this embryonic stage^91^. Another possibility could be that cultured eSSCs lose their ability to support hematopoiesis^92, 93^. We were not able to transplant uncultured eSSCs due to limited number of cells we could obtain in 8 WPC human long bones. Future optimization of the transplantation protocol is needed to further dissect the *in vivo* functions of human and mouse eSSCs. Furthermore, genetic lineage tracing studies would help elucidating the relationship among eSSCs, growth plate SSCs and long bone PSCs in mouse models.

In contrast to endochondral ossification in long bones, intramembranous ossification is the primary way by which calvaria develop^31^. We found 7 osteogenic subsets in 8 WPC calvaria and predicted two distinct sources of osteoprogenitors: 1) from cranial NC lineage cells and 2) from mesodermal lineage cells^94^. Interestingly, we identified a NCDC subset in calvaria that shared similar phenotypic markers as long bone eSSC (Fig. 6d), which represented a transitional state between migratory NC cells and osteoprogenitors. The fact that FOXP1/2 regulons were highly enriched in both long bone eSSCs and calvarial NCDCs suggested a fundamental role of FOXP1/2 in both endochondral and intramembranous ossification. Consistent with this, mouse Foxp1/2 were detected in skeletal progenitors during craniofacial bone development^95^. Unlike long bone eSSCs, NCDCs do not seem to generate chondrocytes (Fig. 6a and Supplementary information, Fig. S6c), which was characteristic of intramembranous ossification. Future studies are needed to test whether NCDCs are evolutionarily conserved in mouse embryonic calvarium, and to prospectively isolate NCDCs for functional analysis of their stem cell activities. Furthermore, the relationships between embryonic NCDCs and calvarial PSCs in postnatal mice could be addressed by genetic lineage tracing studies^4^.

Given that the skeleton repairs in a way that largely recapitulates embryonic development, the skeletogenic mechanisms we uncovered here might help developing novel cell therapies to promote bone and cartilage regeneration, which could ultimately lead to treatments of skeletal disorders such as non-union fracture, osteoporosis and craniofacial defects.

## Materials and Methods

### Human embryonic sample collection

Healthy human embryonic samples were obtained with elective medical termination of pregnancy in the Academy of Military Medical Sciences (the Fifth Medical Center of the PLA General Hospital). All human studies were conducted in accordance with the official ethical guidelines and protocols approved by the Ethics Committee of the Affiliated Hospital of Academy of Military Medical Sciences (ky-2017-3-5). The written informed consent was obtained from all participants before sample collection. Days post fertilization (dpf) of embryos were determined according to the measurement of crown-rump length (CRL) and number of somite pairs, and staged into 5 and 8 weeks post conception (WPC)^96^. The gender of embryos used for scRNA-seq was identified based on the expression of XIST (female) and RPS4Y1 (male)^97^. Sample information was summarized in Supplementary information, Fig. S1a and 6a. The morphology of the embryonic limb bud and long bone was assessed by Hematoxylin-Eosin Staining Kit (Fig. 1a).

### Mice

NOG (NOD.Cg-Prkdc^scid^Il2rg^tm1Sug^/JicCrl) immunodeficient mice (Beijing Vital River Laboratory Animal Technology Co., Ltd.) were used as recipients for renal subcapsular transplantation of human eSSCs. All procedures and protocols were approved by the Ethics Committee of the Academy of Military Medical Sciences (the Fifth Medical Center of the PLA General Hospital).

### Preparation of single-cell suspensions from human limb buds and long bones

Human embryonic limb buds were isolated and transferred to IMDM medium (Gibco) containing 10% fetal bovine serum (FBS) (HyClone) on ice. The tissues were washed with phosphate-buffered saline (PBS) and transferred to pre-warmed digestion medium containing 0.1 g/mL Collagenase I (Sigma) and 0.1 g/mL Collagenase II (Sigma). After vigorous shaking, the samples were incubated at 37 °C for 30 min with gentle shaking every 5 mins. Digestion was terminated by adding IMDM medium containing 10% FBS. After centrifugation at 350 g for 6 min, collected cells were resuspended in FACS sorting buffer (1 × PBS with 1% BSA) for subsequent staining. For long bone specimens, forelimbs and hindlimbs were dissected to obtain humeri, ulnae, radii, femurs, tibiae and fibulae. For calvarial bone specimens, frontal bones, parietal bones and occipital bones were dissected. After cutting by scissors, the long bones or calvarial bones were enzymatically digested as described above. The digested tissues were filtrated with 40 μm strainer to remove cartilage or bone chips, after which cells were centrifugated and resuspended in FACS sorting buffer. The viability of cells was 80-90% by trypan blue staining (0.4%) and 70-80% by 7-AAD staining.

### Flow cytometry

The following antibodies were used: CD45-APC-Cy7 (BD, 557833, 1:50), CD31-Biotin (eBioscience, 13-0319-82, 1:50), Steptavidin-APC-eFlour780 (eBioscience, 47-4317-82, 1:100), CD235a-APC-Cy7 (Biolegend, 349116, 1:50), CD140a-BB515 (BD, 564594, 1:50), PDPN-APC (eBioscience, 17-9381-41, 1:50) and CADM1-PE (MBL, CM004-5, 1:50). Cells were stained in sorting buffer (PBS+1% BSA) for 30 min at 4 °C, washed once and resuspended in sorting buffer with 7-AAD (eBioscience, 00-6993-50, 1:50) as live cell dye. Flow cytometry was performed on BD FACS Aria II. Pre-gating was first done for live cells based on 7-AAD staining. Gating strategies were based on Fluorescence Minus One (FMO) controls. FlowJo v10 software was used for analyzing the flow cytometry data.

### CFU-F culture and mesenchymal sphere assay

For CFU-F cultures, sorted cells were seeded in 6-well plate (4-5 × 10^3^ cells/well) containing culture medium (α-MEM supplemented with 10% FBS, 1% Penicillin/Streptomycin solution and 1 ng/mL bFGF) and incubated at 37 °C with 5% CO_2_. Half of the medium was changed every 3-4 days. At day 10, cells were fixed and stained with crystal violet staining solution. Adherent colonies with more than 50 cells were quantified. Serial CFU-F colony formation was performed by seeding sorted cells in culture medium at clonal density, and serially passaged to generate the secondary and tertiary colonies. For mesenchymal sphere assay, 4-5 × 10^3^ sorted cells were plated in a 6-well ultra-low adherent dish with culture medium and left undisturbed for a week^98^. Half of the medium was changed every week, and the spheres were quantified at day 10.

### Adipogenic, osteogenic and chondrogenic differentiation assays

For nonclonal adipogenic and osteogenic differentiation, sorted cells were cultured for 10 days and replated at a density of 2.0 × 10^4^/cm^2^. Adipogenic differentiation was performed in DMEM (Gibco) supplemented with 10% FBS, 1% Penicillin/Streptomycin, 0.5 μM isobutylmethylxanthine (Sigma), 60 μM indomethacin (Sigma, 17378), 5 μg/ml insulin (Sigma) and 1 μM dexamethasone (Sigma, D2915) for 1 week (medium was changed every 3 days), and quantified by oil red O staining (Sigma) and qPCR. Osteogenic differentiation was performed in osteogenic medium (Cyagen, GUXMX-90021) for 3 weeks (medium was changed every 3 days) and quantified by alizarin red staining (Sigma) and qPCR. The osteogenic differentiation medium contained α-MEM supplemented with 10% fetal bovine serum, 1% Penicillin/Streptomycin, 1% glutamine, 50 μg /ml L-ascorbate acid, 10 mM β-glycerophosphate and 100 nM dexamethasone. For nonclonal chondrogenic differentiation, 2.5 x 10^5^ cultured cells were centrifugated at 1,100 rpm in 15 ml polypropylene conical tubes to form pellets and cultured in chondrogenic medium for 3-4 weeks (medium was changed every 3 days). The chondrogenic medium contained high glucose DMEM (Corning) supplemented with 10 ng/ml TGFβ3 (Peprotech), 100 nM dexamethasone (Sigma), 50 μg/ml ascorbic acid-2-phosphate (Sigma), 1 mM sodium pyruvate (Gibco), 40 μg/ml proline (Sigma) and 1X ITS cell culture supplement (Cyagen) containing 6.25 μg/ml bovine insulin, 6.25 μg/ml transferrin, 6.25 μg/ml selenous acid, 5.33 μg/ml linoleic acid and 1.25 mg/ml BSA. Chondrogenic differentiation was quantified by cryosection of the cell pellets followed by toluidine blue staining and qPCR. For clonal trilineage differentiation, single cells were flow cytometrically sorted into 96-well plates to form single CFU-F colonies. Clonally expanded cells were split into three parts and allowed to differentiate in osteogenic, adipogenic and chondrogenic mediums as described above. Clonal chondrogenic differentiation was also validated by alcian blue and safranin O staining.

### RNA extraction and quantitative real-time PCR (qPCR)

Total RNA was extracted from cells using Trizol reagent (Invitrogen) according to the manufacturer’s instructions. cDNA was prepared using Transgene reverse transcription kit (Transgene). qPCR reactions were prepared using SYBR Green Master Mix (Applied Biosystem) and run on a 7500 Real-Time PCR Systems (Applied Biosystems). A list of the primers used was provided in Supplementary information, Table S5. Human *GADPH* was used as loading control and the relative mRNA abundance was calculated using a comparative CT method.

### Renal subcapsular transplantation

The eSSCs were sorted by flow cytometry and cultured for 7-10 days as previously described^99^. Briefly, 5 ×10^5^ cells were resuspended in 5 μl of Matrigel (BD) on ice and then aspirated into a micropipette (Drummond Scientific, 5-000-2010). A small incision was made near the kidney pole to separate the capsule from the renal parenchyma. Matrigel with cells were injected into the kidney pocket. Eight weeks after transplantation, grafts were dissected and fixed in 4% paraformaldehyde at 4 °C for 12 h, decalcified in 10% EDTA at room temperature for 3 days and then dehydrated in 30% sucrose at 4 °C overnight. Grafts were then cryosectioned at 10 μm and stained by Movat Pentachrome Staining Kit (ScyTek, MPS-1) to demonstrate bone and cartilage differentiation. Immunostaining of collagen I and II were also performed on adjacent sections (see below).

### Immunofluorescent staining

Slides containing renal subcapsular graft cryosections were blocked (10% horse serum and 0.1% Triton-X100 in PBS) at room temperature for 1h and stained with anti-collagen I (Abcam, ab34710, 1:500) and anti-collagen II (Abcam, ab185430, 1:500) antibodies at 4 °C overnight. After washing in PBS (3 × 10 minutes), anti-Rabbit Alexa Fluor 555 (Invitrogen, A31572, 1:500) and anti-Mouse Alexa Fluor 647 (Invitrogen, A31571, 1:500) secondary antibodies were incubated for 1h at room temperature. After washing in PBS (3 × 10 minutes), slides were mounted with ProLong™ Gold Antifade Mountant with DAPI (Invitrogen, P36931). For long bone cryosection staining, the following antibodies were used: anti-PDPN (eBioscience, 17-9381-41, 1:50), anti-CADM1 (abcam, ab3910, 1:100), anti-Rabbit Alexa Fluor 555 (Invitrogen, A31572,1:500) and anti-Rat Alexa Fluor 647 (Invitrogen, A21472, 1:500). Images were acquired with Olympus fluorescence inverted microscope (IX73) and analyzed by ImageJ software.

### Single-cell RNA-sequencing

Samples from different stages were harvested and live cells were sorted based on 7-AAD staining (90-95% viability after sorting). Cells were resuspended at 1 × 10^3^ cells/ml and loaded on Chromium Controller to obtain single cells (10X Genomics). For scRNA-seq libraries construction, Chromium Single cell 3’ Library and Gel Bead Kit V2 (10X Genomics, PN120237) was used to generate single cell gel beads in emulsion (GEM). The captured cells were lysed, and the released RNA were reverse-transcribed with primers containing poly-T, barcode, unique molecular Identifiers (UMIs) and read 1 primer sequence in GEMs. Barcoded cDNA was purified and amplified by PCR. The adaptor ligation reaction was performed to add sample index and read 2 primer sequence. After quality control, the libraries were sequenced on Illumina Hiseq X Ten platform in 150 bp pair-ended manner (Berry Genomics Corporation, Beijing, China).

### Processing of scRNA-seq data

Sequencing data from 10X Genomics were processed with *CellRanger* (version 3.0.1) for demultiplexing, barcode processing and single-cell 3’ gene counting. Human genome reference (GRCh38) was used for sequence alignment. Only confidently mapped, non-PCR duplicates with valid barcodes and UMIs were used to generate the gene-barcode matrix. For quality control, only cells with more than 1, 000 genes and less than 10% of mitochondrial gene were retained for downstream analysis. Cell doublets were removed using *Scrublet* software implemented in python^100^ (https://github.com/AllonKleinLab/scrublet). Briefly, we computed doublet score for each cell by applying *Scrublet* function to each 10X dataset. Then we estimated the number of expected doublets (*N*) with multiplet rates (based on the number of cells recovered) provided by 10X Genomics guideline. Top *N* of cells ranked by doublet scores were determined as doublets (Supplementary information, Fig. S1a and S6a). To correct batch effects among different samples, we applied canonical correlation analysis (CCA) method implemented in Seurat for dataset integration^45^. The union of the top 2,000 genes with the highest dispersion for each dataset was taken to identify anchors using the *FindIntegrationAnchors* function and calculate 30 dimensionalities. We then applied *IntegrateData* function to generate integrated expression matrix, which was used for dimensionality reduction and clustering subsequently. To exclude karyotype abnormalities in human embryos, we applied CNV estimation for single cells in 10X datasets from a previous study^44^. Briefly, we downloaded the expression matrix of non-malignant cells (T cells) and malignant cells as reference cells for the estimation of CNVs. We sampled 100 cells for each 10X dataset and combined them with reference cells to calculate initial CNVs and final CNVs. The CNV correlation score of each single-cell was computed and visualized by heatmap (Supplementary information, Fig. S1b).

### Dimensionality reduction and clustering

To reduce the variation in cell proliferation status that might interfere with single cell analysis, we used the previously reported G1/S and G2/M phase-specific genes to compute scores of S phase and G2M phase, as well as estimate cell-cycle status^101^. We scaled the integrated data with regressing the *S.Score* and *G2M.Score*, and calculated the top 30 principal components (PCs). For dimensionality reduction, we performed Uniform Manifold Approximation and Projection (UMAP) on whole datasets, and used Diffusion map and PCA to visualize the subset of datasets (Supplementary information, Table S3). t-Distributed Stochastic Neighbor Embedding (t-SNE) was applied to visualize the relationships between cell clusters at pseudo-bulk level. For clustering, improved graph-based clustering of the integrated dataset was performed using louvain algorithm after constructing the Shared Nearest Neighbor (SNN) graph. The resolution parameters were set to 0.2 (Supplementary information, Table S3). To ensure the robustness of clustering, we randomly subsampled 1,000 cells from each dataset, and re-processed as previous steps and parameters. The newly identified clusters showed an average assignment of 80% to clusters identified in the whole dataset.

### Species comparative analysis

For comparative analysis between human and mouse datasets, the expression data matrix of mouse E11.5 and E15.5 from GSE142425 were collected^68^. To ensure the comparability, the stage correspondences were identified^102^ and the mouse datasets were processed by the same steps as human datasets, including dimension reduction and clustering. *SciBet* R package (version 1.0)^69^ was used to compare cell subsets identified in limb buds and long bones. We used the expression matrix of human cells as reference dataset and calculated the mean expression values of marker genes across cells with identical cell types. Multinomial models were then built and the query mouse dataset were re-annotated. Sankey plot with *ggalluvial* R package was applied to visualize the matching degree of predicted mouse cell type to the human reference.

### Differential expression analysis

Non-parametric Wilcoxon rank sum test was performed to find DEGs among individual clusters. DEGs were filtered by fold change of more than 2 and cell fraction of more than 20%. DEGs with *P* value adjusted by *benjamini-hochberg* less than 0.01 were considered to be significant (Supplementary information, Table S1).

### Single-cell regulatory network analysis

The analysis of single-cell gene regulatory network was performed using the *SCENIC* package^64^. A standard pipeline implemented in R can be found in https://github.com/aertslab/SCENIC. The expression matrix was loaded onto *GENIE3* for building the initial co-expression gene regulatory networks (*GRN*). The regulon data was then analyzed using the *RcisTarget* package to create TF motifs using hg19-tss-centered-10kb (for human) and mm9-tss-centered-10kb (for mouse) database. The regulon activity scores were calculated with Area Under the Curve (AUC) by the *AUCell* package. Significant regulons enriched in different clusters were calculated by two-sided unpaired t test implemented in Limma R package (version 3.38.3) (Supplementary information, Table S2). The mean regulon activity scores for each cluster were calculated and visualized by heatmap. Predicted target genes of regulon were ranked by *Genie3Weight* value and filtered by normalized enrichment score (NES) of binding motifs (greater than 3). The transcriptional network of TF and predicted target genes was visualized by *Cytoscape* (version 3.6). Edges indicated the *Genie3Weights* and Node size indicated the number of motifs.

### Reconstructing single cell trajectory

Single cell trajectory was analyzed by R package *Slingshot* (version 1.0.0), which infers trajectory by fitting principal curves based on given cell embeddings^81^. After specifying the start or end cluster of the trajectory, cells were projected onto the curve to assign their developmental pseudotime. Specifically, we computed the diffusion map embeddings of OCPs, eSSCs, osteoprogenitors and two subsets of chondrocytes to infer osteo-chondrogenic trajectory. The diffusion components 1 and 3 were used as the input to *Slingshot* (Fig. 3d), and OCP was set as start cluster. For calvarial osteogenesis trajectory, we re-computed the UMAP embedding of NCs, mig_NCs, NCDCs, osteoprogenitors and two subsets of PMSCs, and used UMAP component 1 and 2 as the input to *Slingshot*. The osteoprogenitor subset was set as end cluster (Fig. 6e). To investigate temporally expressed genes changing in a continuous manner over pseudotime, *GAM* function implemented in gam R package was used to find pattern genes along the trajectories. For identification of major patterns, top 200 genes with the most significant time-dependent model fit were retained, and expressions of these genes were smoothed over 20 adjacent cells. To quantify the connectivity of clusters within single-cell graph, the partition-based graph abstraction (*PAGA*) method implemented in Scanpy (version 1.4.3)^103^ was used to generate the abstracted graph.

### RNA velocity

*RNA velocity*^58^ was used for pseudo-time analysis in the integrated dataset of limb buds and long bones (Fig. 1f), as well as OCLC subsets (Fig. 3c). The spliced and un-spliced reads were quantified by the *velocyto* (version 0.17.11) python package with human genome reference. The output loom file was analyzed for velocities of each gene following the pipeline of *scvelo* python package (version 0.1.25)^104^. Count matrix were filtered by top 2,000 highly variable genes and first- and second-order moments were computed for each cell with nearest neighbor set to 30.

### Transcript-averaged cell scoring (TACS)

We adopted TACS as previously described to evaluate cell distribution along selected query genes^73^. For each cell, average expression of the top 100 correlated genes was set as the expression score of the query gene. *Stat_density2d* function implemented in *ggplot2* package was used for visualization. Threshold for partitioning was set to zero.

### Gene functional annotation analysis

Gene ontology (GO) enrichment analysis was performed for DEGs using *clusterProfiler* package^105^. The significant DEGs were used as input to *compareCluster* function and ontology was set to the BP (biological process). The *P* values of enriched GO terms were adjusted by *Benjamini-Hochberg* method and terms were filtered by setting *pvalueCutoff* to 0.05. *Simplify* function was performed to remove redundancy of the enriched GO terms.

### Gene set analysis

GSVA was performed using the *GSVA* R package (version 1.30.0)^106^. We selected gene sets of curated signaling pathways from the MSigDB Database (v7.0, https://www.gsea-msigdb.org) to identify pathways enriched in different limb mesenchymal subsets. The gene-by-cell matrix was converted to gene-set-by-cell matrix and GSVA scores were computed for gene sets with a minimum of 5 detected genes. Significant pathways enriched in different clusters were calculated by two-sided unpaired *t* test implemented in *Limma* R package (version 3.38.3).

### Surface markers and TFs

Surface marker and transcription factor lists were downloaded from the *in silico* human surfaceome (http://wlab.ethz.ch/surfaceome/)^107^ and HumanTFDB3.0 (http://bioinfo.life.hust.edu.cn/HumanTFDB/) database websites (Supplementary information, Table S4).

### Statistics and reproducibility

Values in dot plots were presented as mean ± SEM. Statistical analyses were performed using R and SPSS. The statistical significance of differences was determined using one-way ANOVA with for multiple comparisons. Wilcoxon signed rank test was used to determine the statistical significance of differences for gene expression (2^−ΔΔCt^) analyses (Fig. 4e, 5c and Supplementary information, Fig. S4d, S5d). For single-cell RNA sequencing, three biological replicates for limb bud at 5 WPC and long bone at 8 WPC, and two biological replicates for calvaria at 8 WPC. Clustering for single-cell data were confirmed using subsampling and re-clustering and similar results were obtained as described above. FACS assays were performed at three independent samples for sorting strategies (Fig. 4c). H&E staining and immunostaining were performed at two independent samples (Fig. 1a, 4b and Supplementary information, Fig. S6d). Clonal and Nonclonal differentiation experiments, qPCR assays, renal subcapsular transplantation were performed at three independent samples (Fig. 5a,b,c,d and Supplementary information, Fig. S5b-d).

## Data availability

The accession number for the human scRNA-seq data reported in this paper is GEO: GSE143753. All other relevant data are available from the corresponding authors upon request. The accession number for the count matrices of mouse datasets used in this paper is GSE142425^68^.

---

**Supplementary Figure 1.**
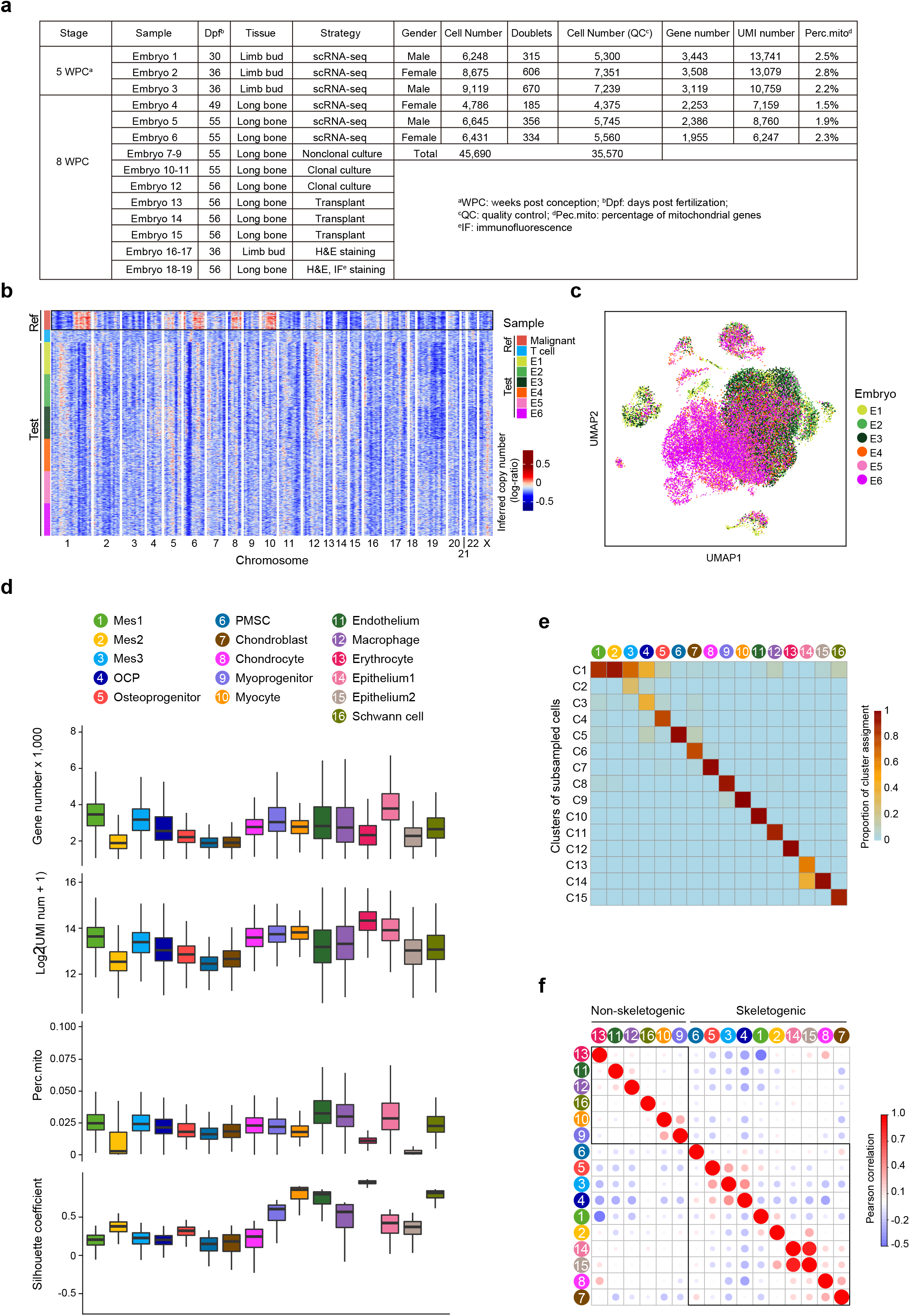
Sample information and data quality control **a,** Table summary of human embryonic limb bud and long bone samples and detailed scRNA-seq information. **b,** CNV scores inferred from transcriptomes of tumor cells, normal T cells (reference cell type) and 100 randomly selected cells from the 6 embryos analyzed by scRNA-seq (test cells). Red: amplifications; Blue: deletions. c, UMAP visualization of the 6 embryos analyzed by scRNA-seq. These included 5 WPC limb buds (E1-3) and 8 WPC long bones (E4-6). **d,** Boxplot showing the number of detected genes, log-transformed UMI counts, percentage of mitochondrial genes and Silhouette coefficient for each subset. **e,** Assessment of the 15 clusters from 6,000 randomly subsampled cells (1000 cells from each embryo) to the 16 subsets annotated in Fig. 1c. **f,** Pearson correlation analysis showing the relationship among the 16 subsets. Hierarchical clustering according to Pearson correlation distinguished skeletogenic (clusters 1-8, 14, 15) and non-skeletogenic subsets (clusters 9-13 and 16).

**Supplementary Figure 2.**
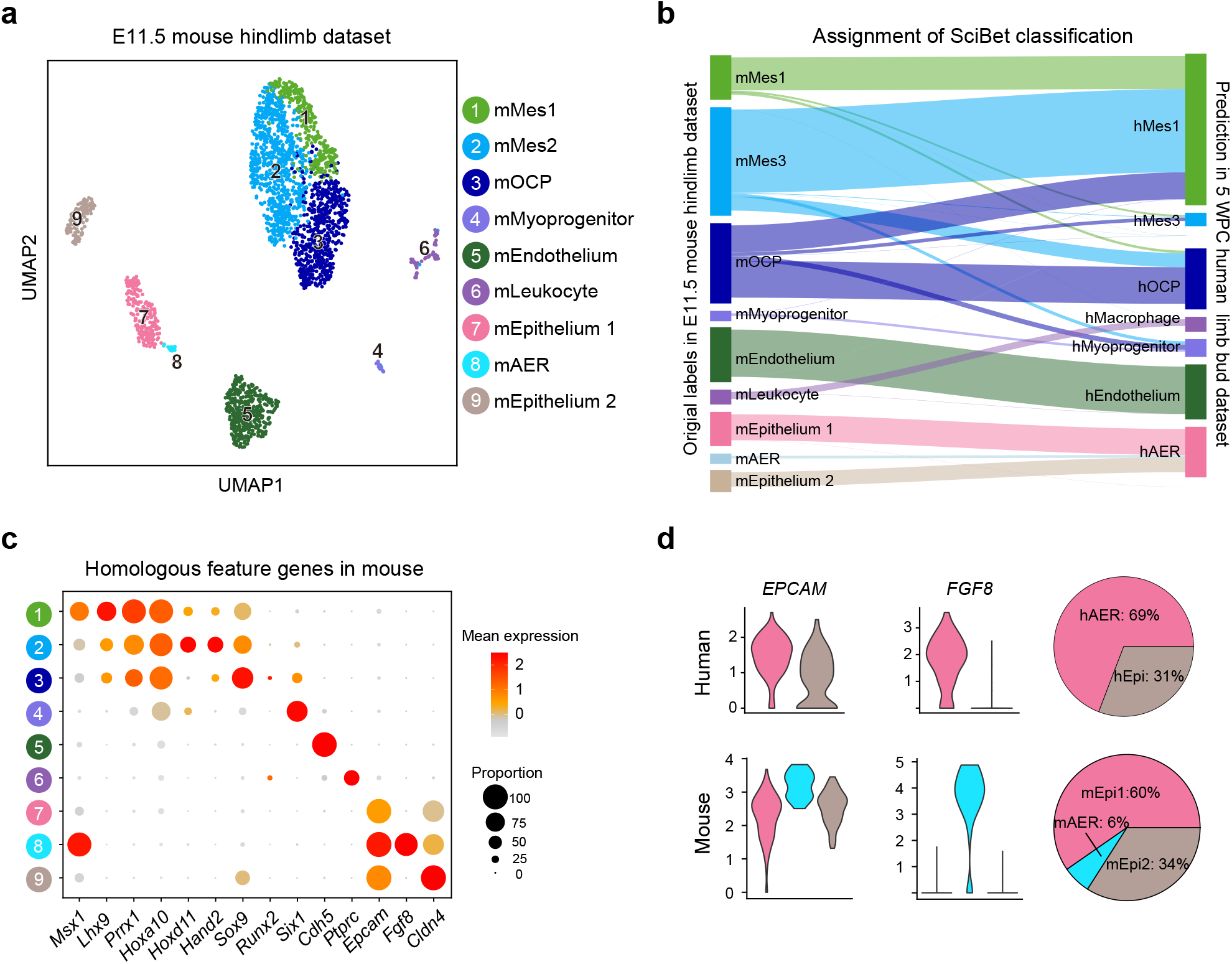
Characterization of E11.5 mouse hindlimb bud mesenchyme and epithelium. **a,** UMAP visualization of 9 cell subsets in E11.5 mouse hindlimb bud dataset. Expression matrix was re-processed and cells were clustered according to the expression of homologous feature genes in human limb bud. **b,** Sankey diagram for assigning E11.5 mouse hindlimb bud subsets to 5 WPC human limb bud datasets. **c,** Dot plots of mean expression of homologous feature genes in E11.5 mouse hindlimb bud subsets. **d,** Violin plots (left) showing the gene expression of EPCAM and FGF8 in human and mouse epithelial subsets. Pie charts (right) showing the proportions of each epithelial subset.

**Supplementary Figure 3.**
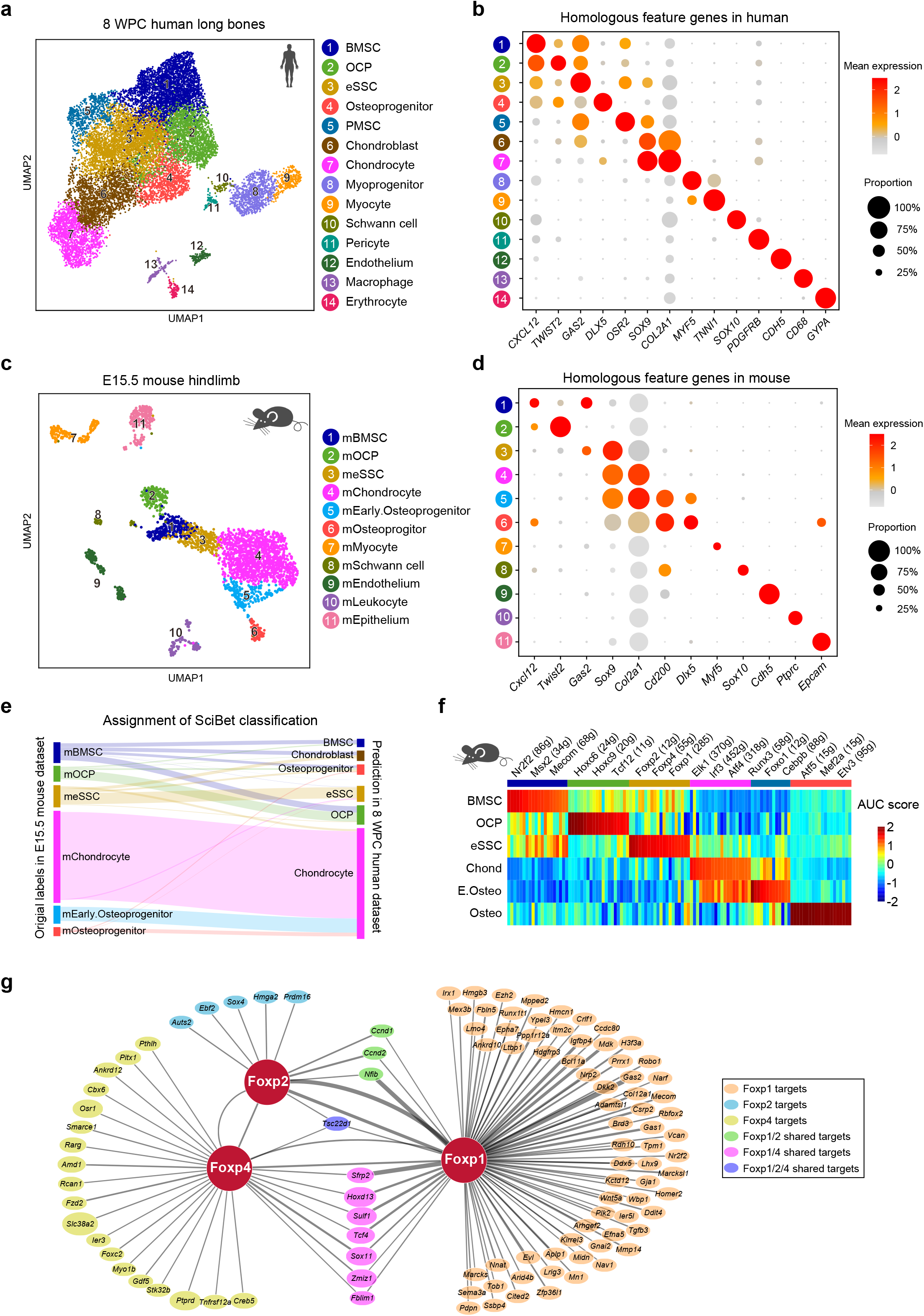
Cross-species comparison between human and mouse embryonic ones during POC formation. **a,** UMAP plot of the 14 subsets in 8 WPC human long bones. **b,** Dot plots showing the expression of human homologous feature genes in the 14 subsets indicated in **(a)**. **c,** UMAP plot of the 11 subsets in re-processed E15.5 mouse hindlimb dataset. **d,** Dot plots showing the expression of mouse homologous feature genes in the 11 mouse hind-limb subsets indicated in **(c)**. **e,** Sankey diagram for assigning mouse E15.5 hindlimb datasets to human 8 WPC long bone datasets. **f,** Heatmap showing the AUC scores of regulons enriched in mouse OCLC subsets. Z-score (column scaling) was calculated. Representative regulons were shown on the right. **g,** The Foxp1/2/4 regulon networks in mouse OCLC subsets. Lines thickness indicated the level of GENIE3 weights. Dot size indicated the number of enriched TF motifs.

**Supplementary Figure 4.**
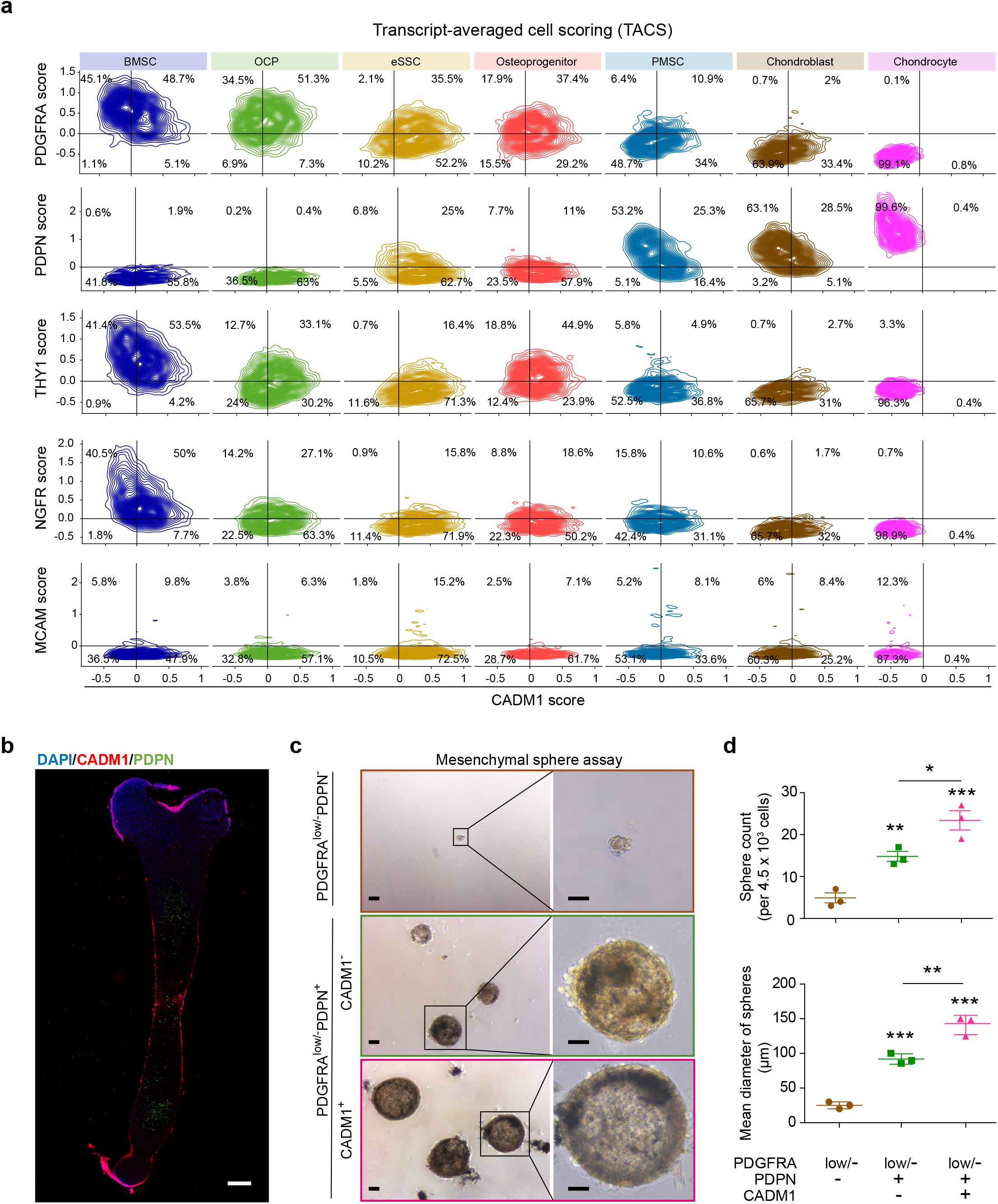
Further in silico and functional analyses of eSSCs. **a,** TACS plots showing the distribution of each OCLC subset between indicated surface marker pairs. Contours outlined regions of increasingly higher cell density. Cell frequencies were shown on the plots. **b,** Representative immunofluorescent image of 8 WPC human femur section stained with DAPI (blue), CADM1 (red) and PDPN (green). **c,** Representative images showing the mesenchymal spheres formed by the 3 populations sorted as in Fig. 4c (left), with magnified views (right). Scale bars: 25 μm. d, Quantification of the number (top) and mean diameter (bottom) of mesenchymal spheres. The statistical significance of differences was determined using one-way ANOVA with multiple comparison tests (LSD). * *P* < 0.05; ** *P* < 0.01; *** *P* < 0.001. Error bars indicated SEM.

**Supplementary Figure 5.**
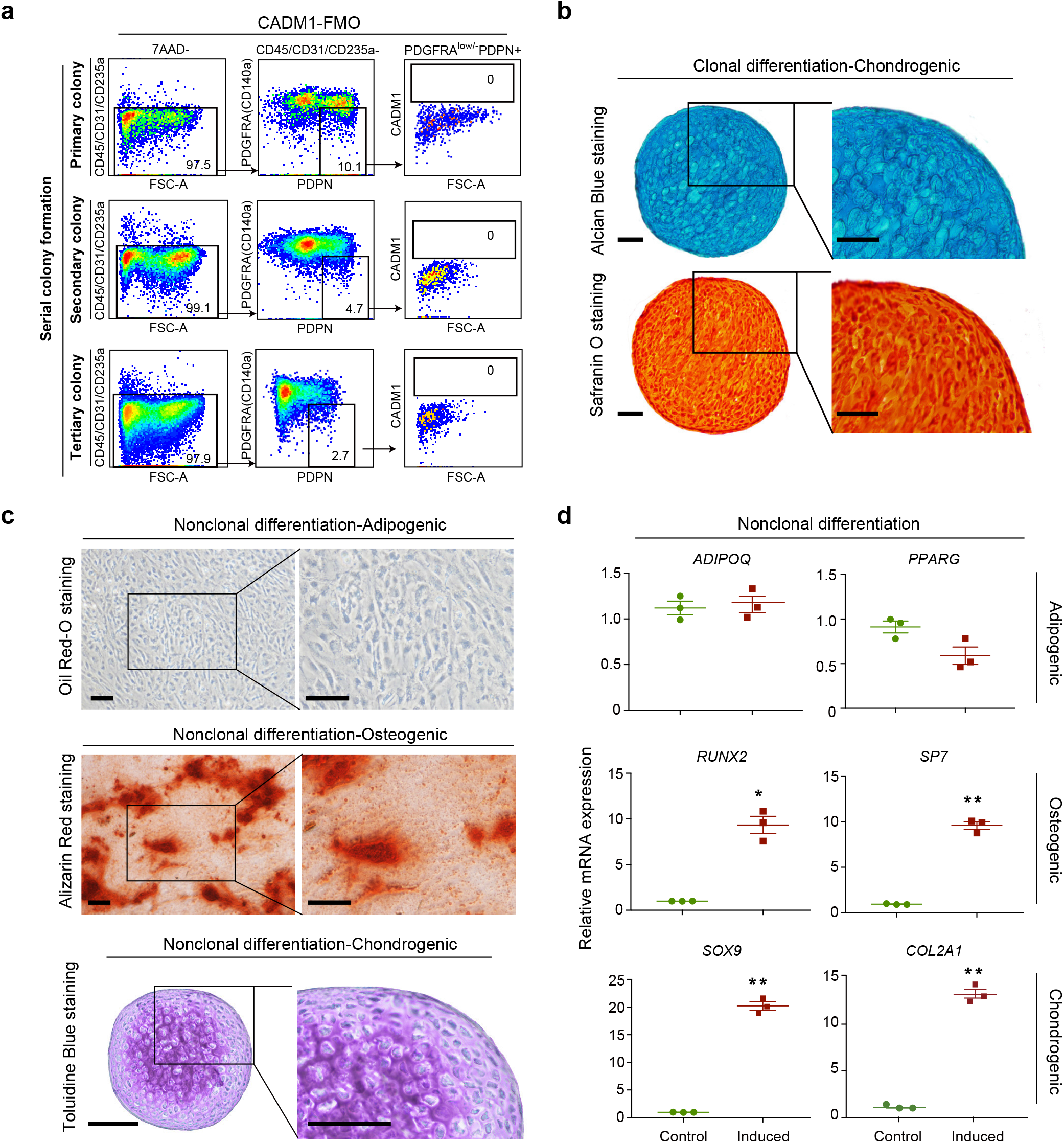
FMO controls and in vitro differentiation of eSSCs. **a,** Fluorescence-minus-one (FMO) controls for eSSC gating strategy in serial colony formation assay (Fig. 5a). **b,** Representative alcian blue (top) and safranin O staining (bottom) after chondrogenic differentiation of clonally expanded eSSCs (PDGFRA low/-PDPN+CADM1+). Magnified images of the boxed areas were shown on the right. Scale bars: 100 μm. **c,** Representative oil red O (top), alizarin red (middle) and toluidine blue (bottom) staining after adipogenic, osteogenic and chondrogenic differentiation of nonclonally expanded eSSCs (PDGFRAlow/-PDPN+CADM1+). Magnified images of the boxed areas were shown on the right. Scale bars: 200 μm. **d,** qPCR analyses of adipogenic, osteogenic and chondrogenic marker genes in nonclonally expanded eSSCs before and after trilineage differentiation in vitro. The statistical significance of differences was determined using Wilcoxon signed rank test. * *P* < 0.05; ** *P* < 0.01. Error bars indicated SEM.

**Supplementary Figure 6.**
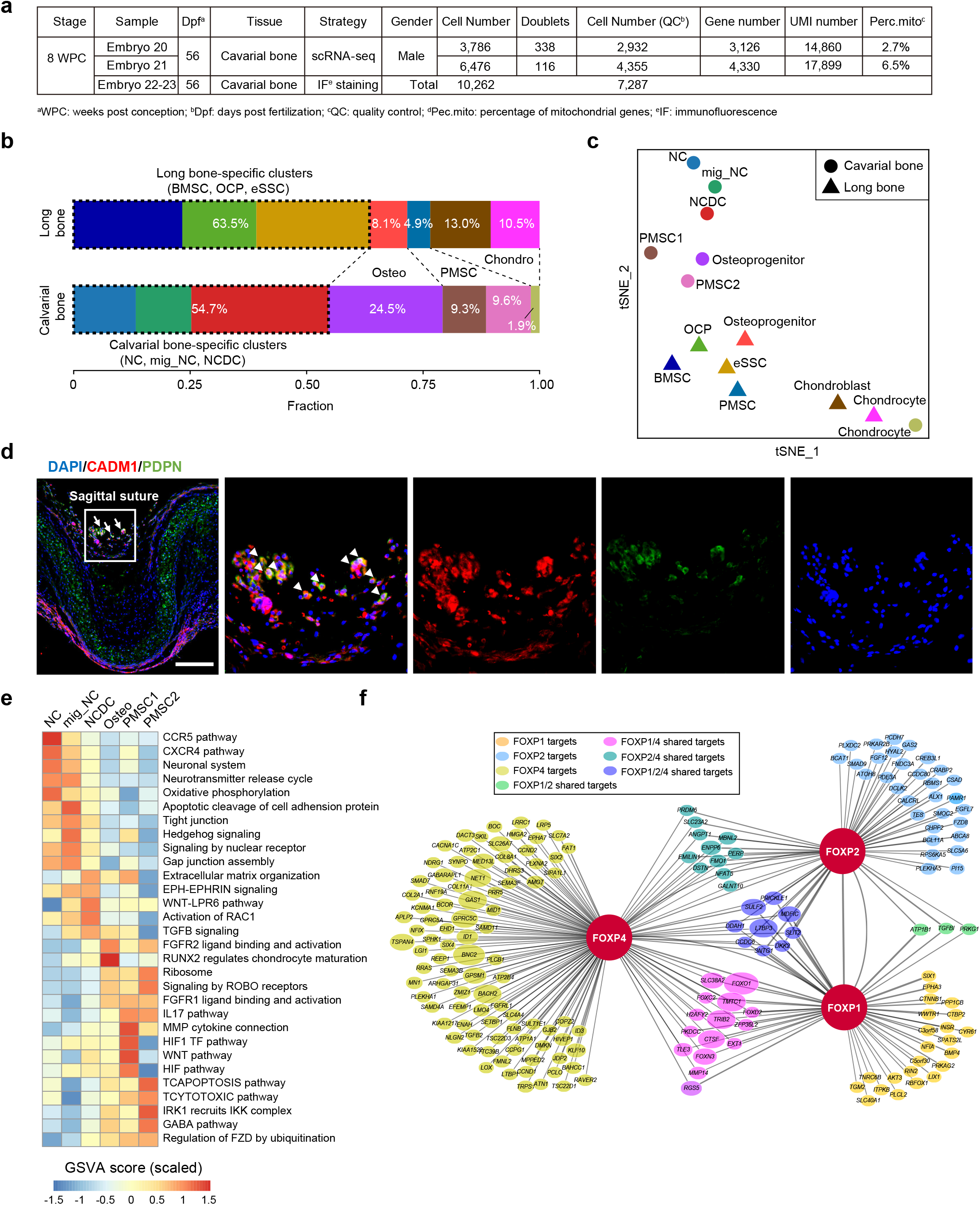
Further characterizations of human embryonic calvaria. **a,** Table summary of the 8 WPC human embryonic calvarial bone samples for scRNA-seq and immunostaining. **b,** Stacked bar charts comparing the distribution of 8 WPC long bone and calvarial subsets. Dashed boxes indicated skeletal site-specific clusters. The three shared clusters (osteoprogenitor, PMSC and chondrocyte) were highlighted by dash lines. **c,** t-distributed stochastic neighbor embedding(t-SNE) projection of indicated subsets from long bones and calvarial bones to compare the transcriptomic similarities at the pseudo-bulk level. **d,** Immunofluorescent images of PDPN+CADM1+ cells in 8 WPC human calvarial bones. Overview of the calvarial region surrounding sagittal suture was shown on the left. PDPN+CADM1+ cells (arrows) were found in the outer layer of sagittal mesenchyme. Arrow heads indicated enlarged PDPN+CADM1+ cells. Merged and single-channel images of DAPI (blue), CADM1 (red) and PDPN (green) were shown. Scale bars: 200 μm. e, Heatmap showing pathways differentially enriched in calvarial bone subsets by GSVA, colored by scaled mean of GSVA scores. f, The FOXP1/2/4 regulon network in 8 WPC human calvarial bone subsets. Line thickness indicated the level of GENIE3 weights. Dot size indicated the number of enriched TF-motif.

## Notes

### Competing Interest Statement

The authors have declared no competing interest.

